# Dyrk1a inhibition with the Novel Compound DYR533: A Cross-Disease Therapeutic Strategy Targeting Amyloidosis, Tau Pathogenesis, and Neuroinflammation

**DOI:** 10.64898/2026.01.20.700091

**Authors:** SK Bartholomew, J Turk, W Winslow, C Foley, A Shaw, S Rokey, S Tallino, MJ Judd, TG Beach, GE Serrano, G Wilms, R Kushwaha, A McMahon, S Ginn, W Becker, C Hulme, T Dunckley, R Velazquez

## Abstract

Alzheimer’s disease (AD) and related dementias are rapidly increasing in prevalence, yet disease-modifying therapies remain largely focused on amyloid-β (Aβ) with limited efficacy against tau pathology and neuroinflammation—key drivers of neurodegeneration and clinical decline. Dual-specificity tyrosine-phosphorylation-regulated kinase 1a (Dyrk1a) phosphorylates tau and amyloid precursor protein and regulates inflammatory signaling, positioning it as a convergence point across pathogenic pathways. We show that brain Dyrk1a protein levels are consistently elevated across ADRDs, replicating findings in AD, confirming prior observations in Pick’s disease, and demonstrating dysregulation in corticobasal degeneration and progressive supranuclear palsy. We developed DYR533, a selective, orally bioavailable, brain-penetrant Type 1 Dyrk1a kinase inhibitor that also inhibits autophosphorylation and reduces kinase abundance. Across three mouse models (3xTg-AD, PS19, Ts65Dn), DYR533 reduced pathological tau hyperphosphorylation, attenuated neuroinflammation, ameliorated amyloidosis, and improved anxiety-like behavior and spatial memory, collectively supporting Dyrk1a inhibition and DYR533 as a therapeutic strategy for ADRD.

## Introduction

The prevalence of neurodegenerative disease continues to increase annually, creating a dire need for therapies that will prevent or slow disease progression. Alzheimer’s Disease and Related Dementias (ADRD) are neurodegenerative disorders that share many of the same pathological features, including aggregation of hyperphosphorylated tau into neurofibrillary tau tangles (NFTs) and neuroinflammation^1–3^; ADRDs include primary tauopathies such as Frontotemporal lobar degeneration – tau dementia (FTLD-tau) as well as Down syndrome (DS), a genetic condition which results in AD in most cases^4^. In DS and AD in particular, an additional hallmark pathology emerges as the amyloid precursor protein (APP) is cleaved by β-secretase (BACE1) forming monomers which aggregate into oligomers and ultimately amyloid-β (Aβ) plaques^5,6^. There have been major advances in therapies for Aβ pathology, mainly based on immunotherapy with monoclonal antibodies^7,8^. However, Aβ therapies have evidenced varied outcomes, and their use is limited to dementias with Aβ pathology. A multifaceted approach that improves multiple pathologies by targeting a single effector would likely provide more robust effects and could potentially be combined with approved amyloid therapies for increased effects^9^.

A targetable protein that is dysregulated across multiple ADRD is dual-specificity tyrosine phosphorylation-regulated kinase 1a (Dyrk1a). While Dyrk1a is expressed in multiple cell types in the brain and periphery, its highest expression is in neurons and microglia^10^. Prior work has shown that Dyrk1a RNA and protein levels are elevated in post mortem brain tissue of individuals with AD and primary tauopathies, including Pick’s disease^11–14^. In humans with DS, Dyrk1a is triplicated within the DS critical region on chromosome 21 (HSA21) resulting in increased expression^15^. Dyrk1a phosphorylates APP at threonine 668, which is associated with enhanced β-secretase (BACE1) processing and increased production of Aβ peptides^16,17^. Dyrk1a phosphorylates pathologically relevant tau epitopes, such as Threonine 181 (Thr181) and Serine 396 (Ser396), promoting the formation of NFTs^18–20^. Dyrk1a can also prime other kinases that hyperphosphorylate tau, including Glycogen synthase kinase-3β (GSK-3β)^21^. Beyond its effects on tau, Dyrk1a contributes to neuroinflammatory responses, as reducing microglial Dyrk1a has been shown to dampen inflammation following a lipopolysaccharide (LPS) challenge^22,23^. Additionally, individuals with mutations in Dyrk1a present with intellectual disabilities^24^. Increased expression of Dyrk1a induces perturbations in long-term potentiation contributing to cognitive dysfunction^25^. Reducing elevated Dyrk1a protein levels in rodent models of AD, DS, and Dyrk1a disorders has been shown to improve performance on behavioral and cognitive tasks^26–32^. Collectively, these findings support Dyrk1a as a promising therapeutic target for multiple aspects of diverse neurological disorders.

The development of Dyrk1a inhibitors has emerged as an active area in the field^33–35^. Dyrk1a inhibitors naturally occur in the environment and have often been used as starting points to develop improved novel inhibitors. For example, the Dyrk1a inhibitor, Leucettine 41 (L41) is derived from Leucettamine B produced by the marine sponge *Leucetta microraphis*^36^. Dyrk1a belongs to the CMGC family, which includes cyclin-dependent kinases (**C**DKs), mitogen-activate protein kinases (**M**APKs), glycogen synthase kinases (**G**SKs) and CDK-like kinases (**C**LKs)^37^. Dyrk1a inhibitors can be potent yet promiscuous compounds, often inhibiting multiple CMGC family kinases or other off-target proteins, potentially affecting mechanisms that can lead to adverse outcomes during long-term treatment^35,38,39^. For example, harmine derivatives are potent Dyrk1a inhibitors, but their strong affinity towards monoamine oxidase, make them generally unsuitable for clinical use^40^.

We previously published on a Dyrk1a inhibitor, termed DYR219^41^. Although DYR219 improved AD-like pathology in the 3xTg-AD mouse model, enhancements in half-life, affinity, and selectivity are required to maximize its therapeutic potential. Here, we introduce and test a novel and superior small molecule Dyrk1a inhibitor, termed DYR533, which exhibits superior blood-brain barrier (BBB) penetrance, excellent oral bioavailability, and is highly selective within the kinome. We hypothesized that DYR533 would inhibit the function of the kinase domain of Dyrk1a in the brain and would also reduce Dyrk1a protein levels by preventing its autophosphorylation, in turn reducing amyloidosis, tau hyperphosphorylation and inflammation in multiple mouse models of neurodegenerative diseases. We found that DYR533 reduced Aβ burden in both the 3xTg-AD model of AD and Ts65Dn model of DS, while reducing tau and inflammation in these models and in the PS19 model of primary tauopathy. These results establish DYR533 as a potentially valuable, multifaceted therapeutic option that improves pathologies shared across numerous neurodegenerative disorders.

## Methods and Materials

### Human Tissue

Human postmortem superior frontal gyrus (Sfg) tissue was obtained through the Arizona Study of Neurodegenerative Disorders and Brain and Body Donation Program^42,43^. We used 22 sex-balanced samples of Pick’s disease (n = 4), corticobasal degeneration (CBD; n = 3), and progressive supranuclear palsy (PSP; n = 8), as well as healthy controls (HC; n = 7) to assess Dyrk1a protein levels. We also obtained thirty-six sex-balanced samples of blood serum from Control (Con, n = 12, zero to sparse Consortium to Establish a Registry for Alzheimer’s disease (CERAD) neuritic plaque density, Braak neurofibrillary stage ≤ III), AD Moderate (AD Mod; n = 12, moderate to frequent CERAD neuritic plaque density, Braak neurofibrillary stage = IV), and AD Severe (AD Sev; n = 12, frequent CERAD Neuritic plaque density, Braak neurofibrillary stage VI). We also obtained frontal cortical (FC) post-mortem tissue from a subset of these same individuals (Con = 9, AD Mod = 12, AD Sev = 10). AD Mod and Sev corresponded with the National Institute on Aging-Reagan Institute (NIA-RI) intermediate and high classifications, respectively^44^. Pathological assessment of human cases was performed as previously described^42,43^. We used commercially available enzyme-linked immunosorbent assay (ELISA) kits to quantify Dyrk1a protein levels (LSBio, Cat #LS-F8931) in both Sfg and FC tissue, and in blood serum. We also quantified levels of pro-inflammatory cytokine Tumor necrosis factor-⍺ (TNF-⍺; Abcam, Cat #ab181421) in FC tissue and blood serum.

### Animals

3xTg-AD mice (MMRRC Strain #034830-JAX) homozygous for human genes *APP* with Swedish mutation, *Presenilin 1 (PSEN1)* with the M146V mutation, and *microtubule-associated protein tau* (MAPT) with P301L mutation were generated as previously described on a C57BL6/129Svj hybrid background^45,46^. All 3xTg-AD mice and background strain non-transgenic controls (NonTg) were bred in-house (n = 12 - 15/group). Notably, 3xTg-AD males show substantial neuropathological variability, even between littermates, which is not observed in females. Therefore, as in most recent studies using the 3xTg-AD mouse, we only included female mice^47–49^. Two-month-old male and female PS19 mice (Jackson Laboratory Strain #008169) and NonTg littermates (controls) were purchased from Jackson Laboratories (n = 16 - 18/group, balanced for sex). PS19 mice are heterozygous for the human P301S mutation of the *MAPT* gene, a known genetic cause of early onset FTD in humans^50^. Trisomic (3n) Ts65Dn mice (Jackson Laboratory Strain #005252) and disomic (2n) littermates (controls) were also purchased from Jackson Laboratories (n = 15 - 18/group, balanced for sex). 3n mice carry an additional translocation chromosome composed of segments of murine chromosome 16 and 17, resulting in trisomy of approximately two-thirds of orthologous genes on HSA21 including Dyrk1a^51–54^. While Ts65Dn mice do not typically develop Aβ or tau inclusions, they recapitulate the early degeneration of Basal forebrain (BF) cholinergic neurons seen in DS and AD, which is thought to drive cognitive and memory dysfunction^55^. All mice were maintained on a 12-hour light/dark cycle at 23°C with *ad libitum* access to food and water and group-housed with four to five mice per cage. All animal procedures were approved by the Institutional Animal Care and Use Committee of Arizona State University (IACUC). Each animal was weighed weekly, beginning at the start of drug regimen (baseline) until euthanasia. Body weight was used to accurately calculate the amount of drug given to each mouse.

### Dyrk1a tyrosine autophosphorylation assay

To assess Dyrk1a co-translational tyrosine autophosphorylation in the presence of DYR533 and other inhibitors, we conducted an *in vitro* translation assay and quantified phosphotyrosine (pTyr) levels. The NEBExpress® Cell-free E. coli Protein Synthesis System (New England Biolabs, Cat #E5360S) was used to express a Dyrk1a construct comprising the kinase domain (residues 126–490) with an N-terminal His tag. Reactions were run in a total volume of 6.25 μL with 40 ng/μL pEXP17-DYRK1A (kind gift of Ulli Rothweiler, Tromsø) at 21°C for 3 h. The Dyrk1a inhibitors L41 (Adipogen Cat#MR-C0023)^39,56^, DYR533 and CaNDY^57^ were added at 1, 3 and 10 μM concentrations (∼1% final DMSO concentration) and compared to DMSO control. Tyrosine autophosphorylation of Dyrk1a was detected by western blot using a phosphor-HIPK2 (pTyr361) antibody (Thermo Fisher Scientific Cat #PA5-13045, RRID:AB_10987115; 1:500 dilution) as described previously^58^. The total amount of the recombinant Dyrk1a construct was assessed by detection of the His6-Tag (mouse anti-His antibody, Pharmacia GE Healthcare, Cat #27-4710). Band intensities were quantitated using ImageQuant TL Analysis software (GE Healthcare Life Sciences).

### Cell-based Dyrk1a degradation assay

The NEBuilder HiFi DNA Assembly Cloning Kit (New England Biolabs, Cat #E5520S) was used to insert rat Dyrk1a cDNA into a version of pcDNA5/FRT/TO already equipped with a C-terminal HiBiT sequence^58^. The resulting recombinant Dyrk1a-HiBiT construct carries a short N-terminal deletion, because a nonsense mutation was inadvertently introduced at codon 51 during the PCR-based cloning procedure. Stable Flp-In T-Rex HEK293 cell lines for doxycycline-inducible expression of Dyrk1a-HiBiT were established by Flp recombinase-mediated integration of the pcDNA5/FRT/TO expression vector according to the manufacturer’s instructions (Invitrogen, Cat#V601020). Cells were grown in Dulbecco’s modified Eagle medium/F-12 (DMEM/F-12, Thermo Fisher Scientific, Cat#11330057) supplemented with 10% fetal bovine serum and maintained at 37°C in a humidified 5% CO_2_ atmosphere. For the degradation assays, cells were resuspended in cell culture medium containing 2 µg/mL doxycycline and dispensed into 96-well tissue culture plates (Sarstedt, Cat#83.1835) at a density of 10,000 cells per well in a volume of 90 µL. The following day, compounds were added to the wells (in triplicates) by dispensing 10 µL of 10-fold concentrated working solution to achieve the desired final concentrations of the compounds or the solvent control (0.1% DMSO). After 24 h of treatment, the culture medium was carefully removed and cells were lysed in 100 µL of ice-cold Passive Lysis Buffer (Promega Corporation, Cat#E1941) on an orbital shaker for 30 min. Cell lysates were cleared by centrifugation (20 min at 20,000 x g), and aliquots of 10 µL of the supernatants were used for Nano-Glo HiBiT Lytic Assay (Promega Corporation, Cat#N3030) to determine the concentration of HiBiT-tagged proteins in the soluble fraction.

### Dyrk1a Inhibitor (DYR533) and Dosing

DYR533 was solubilized in a solution of equal parts polyethylene glycol (molecular weight 400; PEG400) and 0.9% Sodium chloride (NaCl). The PEG400 and 0.9% NaCl solution (1:1) without DYR533 served as the vehicle (Veh). To determine the selectivity of DYR533 a KINOMEscan (*Eurofins*) was performed. This widely used competitive binding assay screens for kinase inhibitors by assessing the ability of the compound of interest to compete with an immobilized ligand for binding to a DNA tagged kinase. Quantification was performed using qRT-PCR and reported as an S(35) score. The experimental conditions used in the KinomeProfiler-Eurofins/CEREP radiometric protein kinase assays (a total of 403 kinases) are described in detail at www.eurofinsdiscovery.com/solution/kinase-profiler.

Female 3xTg-AD and NonTg mice were randomly assigned to one of four groups starting at seven and a half months of age: Veh, 1.0 mg/kg, 2.5 mg/kg, and 5.0 mg/kg of body weight. Daily intraperitoneal (IP) injections were administered for approximately two months. Dosing of 3xTg-AD mice began prior to the onset of widespread tau pathology^59^. Male and female PS19 and NonTg littermates were randomly assigned to one of four dosing regimens starting at four months of age: Veh, 1.0 mg/kg, 2.5 mg/kg, and 5.0 mg/kg of body weight. Daily IP injections were carried out for four months, beginning prior to the onset of widespread tau pathology in PS19 mice^50^. 3n and 2n mice were randomly assigned to one of four dosing regimens: Veh, 0.625 mg/kg, 2.5 mg/kg, and 10 mg/kg of body weight. Daily IP injections began at four and a half months of age, prior to the development of soluble amyloid and tau accumulation and BF degeneration and continued for three and a half months^60^. Doses for each study were determined based on preliminary two-week maximum tolerable dose studies conducted for each mouse strain.

### Behavioral testing

#### Rotarod

PS19 and littermate NonTg mice underwent three days of Rotarod testing (Rota-Rod Advanced, TSE Systems) at seven and a half months of age to assess motor coordination and endurance^61^ as previously described^62,63^. Briefly, each mouse was trained for two days, followed by a probe day, with six time-spaced trials per day. Trials lasted 90 seconds or until the mouse fell off the spinning rod. On training days, the rotation speed of the rod increased by 0.75rpm/s over 20 seconds to a maximum of 15 rpm. On the probe day, the rod accelerated at 1rpm/s from 0 to 60 rpm. The system was connected to computer software (TSE Rotarod, version 5.1.2, TSE Systems) that measured and recorded each mouse’s latency to fall.

#### Morris Water Maze

To assess hippocampal-dependent spatial learning and memory, PS19 mice and NonTg littermates were tested in the Morris Water Maze (MWM) as previously described^63^. The MWM consists of a one-and-a-half-meter diameter pool filled with 23-24°C water, tinted white with non-toxic paint to obscure the view of the platform. A clear platform was submerged one cm below the water in one of the pool’s quadrants, which mice had to locate to be removed from the pool. All mice received four 60s trials per day, with a 30s rest period between each trial for five consecutive days. There were prominent spatial cues placed around the testing room. The hidden platform remained in the same quadrant for all trials and mice, while the start location varied pseudo-randomly across all trials. During each trial, mice were required to locate the hidden platform to escape the pool. If mouse failed to reach the hidden platform in 60s, they were gently guided to its location and remained on the platform for 10s. On the sixth day, a probe trial was conducted in which the platform was removed, and each mouse underwent one 60s trial. All trials were recorded with a video camera, and data was analyzed via EthoVisionXT (Noldus Information Technology).

### Blood Collection and Plasma Extraction

Blood was drawn from 3xTg-AD and PS19 study mice via the submandibular vein at the end of each study. A total of 150-200 μL of blood was collected into EDTA-lined tubes (K_2_EDTA; BD, Cat #365974) and inverted ten times to ensure anticoagulation, as previously described^47,62^. Tubes were kept on ice for 60-90 minutes, then centrifuged at 455 x g for 30 minutes at 4°C to separate phases. The plasma layer was decanted and frozen at −80°C for future analysis.

### Tissue Harvesting and Processing

Mice from the PS19 and Ts65Dn studies were euthanized at eight months and mice from the 3xTg-AD study were euthanized at ten months. Mice were anesthetized with a mixture of Ketamine (120 mg/kg) and Xylazine (6 mg/kg) prior to transcardial perfusions with cold 1x Phosphate-Buffered Saline (PBS). Brains from the 3xTg-AD and PS19 studies were extracted and cut along the midline; one half hemisphere was fixed in 4% paraformaldehyde in 1x PBS for 48 hours, then changed into 0.02% sodium azide for storage until sectioning, and the remaining hemisphere was dissected to isolate the hippocampus (Hp) and cortex (Ctx) and frozen for protein extraction. For the Ts65Dn study, following perfusion, the BF was dissected, as previously described^60,64^ and frozen for protein extraction. Frozen brain samples were homogenized in a tissue protein extraction reagent (TPER, Cat #78510), supplemented with protease (Roches Applied Science, Cat #11836153001) and phosphatase (Millipore, Cat #524625-1SET) inhibitors. Samples were centrifuged for 30 minutes at 21,130 x g and the supernatant was collected as the soluble fraction for ELISAs and BioRad Luminex assay. 70% formic acid was added to the remaining pellet from 3xTg-AD and PS19 study samples which were homogenized and centrifuged as previously mentioned. The insoluble supernatant was collected and added to a neutralization buffer at a 1:9 ratio to be used as the insoluble fraction for ELISAs.

### ELISAs and BioRad Luminex Multi-plex Cytokine Assay

We used commercially-available ELISAs for assessment of homogenates from multiple brain regions and blood plasma. Murine Dyrk1a was quantified (LSBio, Cat #LS-F7258) in soluble Hp and Ctx homogenates for the 3xTg-AD and PS19 studies and in soluble BF homogenates for the Ts65Dn study. For both the 3xTg-AD and PS19 studies, we measured human phosphorylated tau (pTau) at Threonine 217 (T217) (MyBioSource, Cat #MBS1608795) of soluble Ctx only and, pTau T181 (Invitrogen-Thermo Fisher, Cat #KHO0631) and S396 (Invitrogen-Thermo Fisher, Cat #KHB7031) of soluble and insoluble Hp and Ctx. We prioritized the use of Hp and Ctx tissue for T181 and S396 and used the remaining Ctx tissue to quantify T217 and the expression of pro- and anti-inflammatory cytokines and chemokines. For the 3xTg-AD study, we also measured Hp and Ctx soluble and insoluble human Aβ_40_ (Invitrogen-Thermo Fisher, Cat #KHB3481) and Aβ_42_ (Invitrogen-Thermo Fisher, Cat #KHB3441). Murine TNF-⍺ (Abcam, Cat #ab208348) was quantified in the Hp, Ctx and blood plasma from the 3xTg-AD and PS19 studies to assess inflammation. To evaluate neuropathology in the Ts65Dn study, murine pTau at T181 (MyBiosource, Cat #MBS9905654) and murine Aβ_40_ and Aβ_42_ (Invitrogen-Thermo Fisher Scientific, Cat #KMB3481 and #KMB3441, respectively) were measured in soluble BF homogenates, as Ts65Dn mice do not possess human transgenes. Additionally, we used the Bio-plex®-200^65^ with Bioplex Manager software (Version 6.2) to quantify the levels of 23 different pro- and anti-inflammatory cytokines and chemokines (Bio-Plex mouse cytokine 23-plex kit; Bio-Rad, Cat #M60009RDPD) in Ctx tissue from the PS19 mice and BF tissue from the Ts65Dn mice, as previously described^47^. GM-CSF was excluded from both studies because values fell outside of the detectable range. All samples were run in duplicate wells.

### Statistical analysis

Analyses for the 3xTg-AD and PS19 studies were conducted using GraphPad Prism 10.3.0. One-way Analysis of Variance (ANOVA) were conducted to assess statistical significance of human Dyrk1a and TNF-⍺ measures. Correlation analyses were performed between human Dyrk1a Sfg and FC levels and pathological measures, including the last Mini-Mental State Examination (MMSE) score, Braak stage, CERAD neuritic plaque density, and brain weight. For *in vitro* assays, to assess Dyrk1a tyrosine autophosphorylation and degradation, treatment effects were analyzed using repeated measures ANOVA and Dunnett’s multiple comparison test to compare treated samples with the Veh. Weight change, Dyrk1a levels, MWM data (latency to platform, speed, distance, thigmotaxia) and Rotarod training days 1 and 2 were analyzed using two-way ANOVA. Probe day for MWM and Rotarod, pTau, TNF-⍺, Aβ and BioPlex cytokine/chemokine quantifications were analyzed using one-way ANOVA. Post hoc corrections were used as recommended by Prism. For the Ts65Dn study, Multiple Analysis of Variance (MANOVA) in SPSS 29.0.2.0 (20) were used to examine main effects of sex, genotype and dose and interactions of Dyrk1a, pTau T181, and Aβ levels. Analyses of individual cytokines/chemokines from the Bioplex readout were conducted using one-way ANOVA after collapsing sexes, as no significant main effect of sex nor interaction was found. Post-hoc testing was performed in GraphPad Prism 10.3.0 with Šidák’s correction for all, except Eotaxin which violated the assumption of normality. For Eotaxin, a Brown-Forsythe ANOVA with Dunnett’s T3 corrections for multiple comparisons was used.

## Results

### Dyrk1a protein levels are significantly elevated in ADRD and correlate with clinical and neuropathological outcomes

Statistical outputs for all analyses are presented in (Supp. Table 1). Previous reports have shown significant elevations of Dyrk1a in human ADRD^11–14^. Here, we first quantified Dyrk1a protein levels in superior frontal gyrus (Sfg) tissue from individuals with primary tauopathies—including Pick’s, corticobasal degeneration (CBD), and progressive supranuclear palsy (PSP) – and healthy control (HC)^42^ and found significant elevation in all disease cases compared to healthy control (HC; Fig. 1A; *p <* 0.0001). We found a strong negative correlation between Dyrk1a and last MMSE^66^ (Fig. 1B; *p =* 0.0324), and a positive correlation with Braak stage^67^ (Fig. 1C; *p=* 0.0088). We also quantified Dyrk1a in human blood serum (serum-Dyrk1a) and frontal cortex (FC) brain tissue (FC-Dyrk1a) from Control (Con), AD Moderate (AD Mod), and AD Severe (AD Sev) cases and found elevation in individuals with AD (Fig. 1D; *p =* 0.0002), where AD Sev had higher levels than both Con (*p =* 0.0009) and AD Mod (*p =* 0.0006) cases. FC-Dyrk1a protein levels were also significantly elevated across AD disease progression (Fig. 1E; *p <* 0.0001), where AD Sev had higher protein levels compared to both Con (*p <* 0.0001) and AD Mod (*p =* 0.0027); AD Mod showed higher levels than Con (*p =* 0.0193). Additionally, FC-Dyrk1a negatively correlated with last MMSE^66^ (Fig. 1F; *p =* 0.0035), while positively correlating with both CERAD neuritic plaque density^68^ (Fig. 1G; *p <* 0.0001), Braak stage^67^ (Fig. 1H; *p <* 0.0001), and negatively correlating with brain weight (Fig. 1I; *p =* 0.0077).

**Figure 1:**
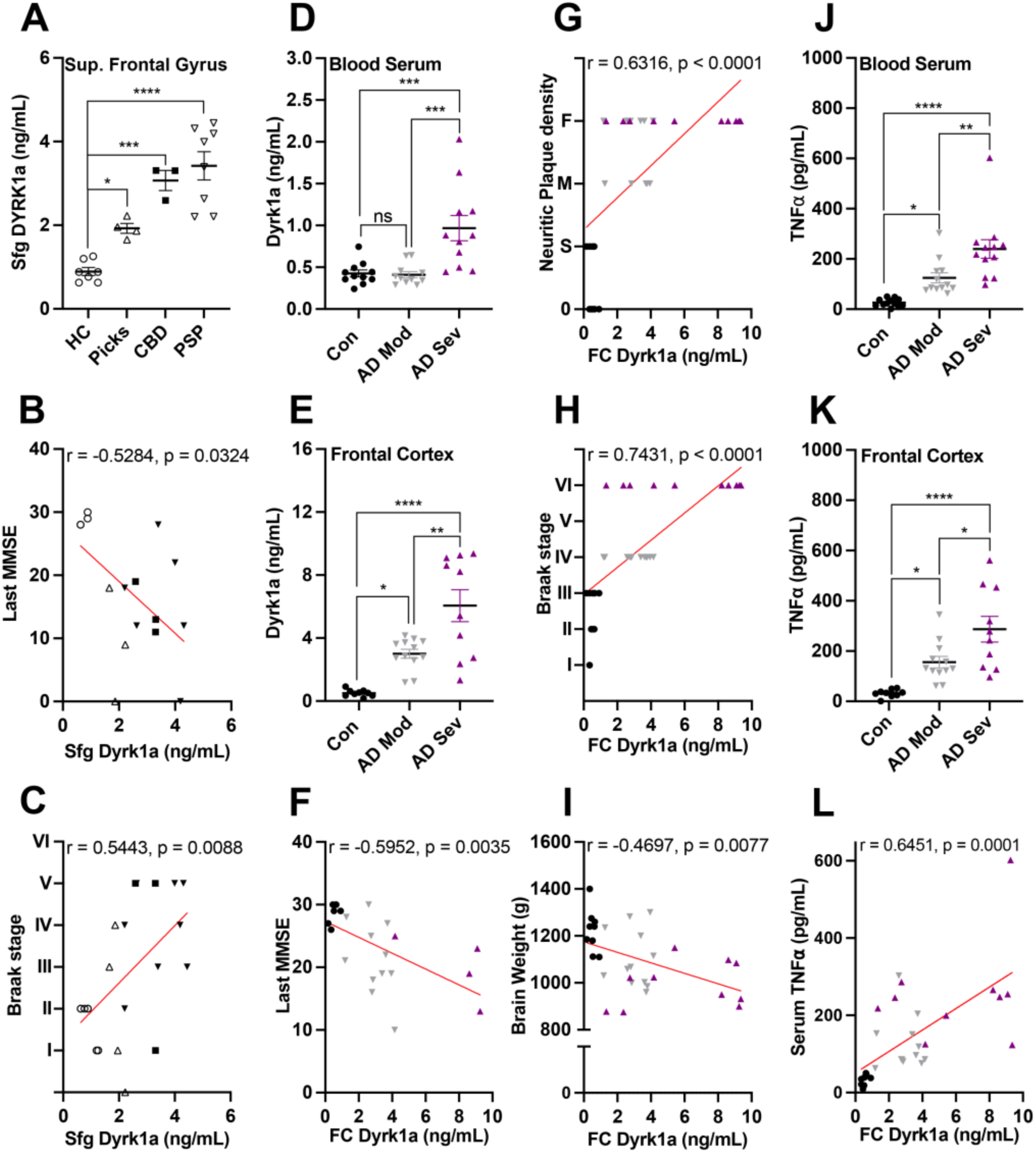
Dyrk1a protein levels are elevated across tauopathies and correlate with clinical, neuropathological and neuroinflammatory factors. (A) Dyrk1a levels were significantly increased in post-mortem superior frontal gyrus (Sfg) tissue from primary tauopathy cases—including Pick’s disease, corticobasal degeneration (CBD), and progressive supranuclear palsy (PSP)—compared with healthy controls (HC). (B–C) Sfg Dyrk1a levels showed significant correlations with Mini-Mental Status Exam (MMSE) scores and Braak stage. (D–E) Dyrk1a protein levels were quantified in post-mortem blood serum and frontal cortical (FC) tissue from Control (Con), AD Moderate (AD Mod), and AD Severe (AD Sev) individuals. (F–I) FC Dyrk1a levels were correlated with clinical and pathological measures across AD progression (CERAD neuritic plaque density: 0 = none, S = sparse, M = moderate, F = frequent). (J–K) TNF-⍺ levels were quantified in blood serum and FC tissue across disease stages. (L) FC Dyrk1a levels correlated with serum TNF-⍺. Data represent mean ± SEM. ns = not significant; *p < 0.05, **p < 0.01, ***p < 0.001, ****p < 0.0001. FC TNF-⍺ also increased with AD progression (Fig. 1K; *p <* 0.0001), where AD Sev had higher levels compared to AD Mod (*p* = 0.0198) and Con (*p* < 0.0001); AD Mod exhibited higher TNF-⍺ compared to Con (*p =* 0.0351). Notably, we found a positive correlation between human blood serum TNF-⍺ and FC-Dyrk1a (Fig. 1L; *p =* 0.0001). Our findings across independent cohorts demonstrate for the first time that Dyrk1a protein levels are elevated in CBD and PSP, extending prior reports of increased Dyrk1a in Pick’s disease^11^ and are consistent with observations of concomitant elevations in Dyrk1a and TNF-α in both peripheral blood and brain tissue across AD progression^11,12^.

Neuroinflammation is another characteristic of ADRD^3,69^. We quantified levels of tumor necrosis factor (TNF)-⍺, which have been shown to contribute to neurodegeneration when systemically elevated^70^. AD cases showed elevated blood serum TNF-⍺ (Fig. 1J; *p <* 0.0001), where AD Sev cases had higher levels compared to AD Mod (*p <* 0.0009) and Con (*p* = 0.0006).

### DYR533 inhibits intramolecular tyrosine autophosphorylation and induces degradation of Dyrk1a *in vitro*

Our Dyrk1a inhibitor, DYR533 (Fig. 2A) was screened for interactions with 403 human WT kinases (the “kinome”) using a commercially available fee-for-service competitive binding assay (KINOMEscan – *Eurofins*) (Fig. 2B)^56^. Larger circles indicate stronger binding, and kinases are grouped by family, with DYRK and CLK kinases circled within the CMGC family. Results from the kinome scan are summarized in an S(35) score, corresponding to the proportion of the total kinases with signal inhibited to 35% of the control value for each kinase^71^ (Fig. 2C). Follow-up for key kinases with significant reduction in activity as shown in the kinome screen (i.e. <35% of control values) were as follows: Dyrk1a (K_d_ 1.4nM), Dyrk1b (K_d_ 11nM), Dyrk2 (K_d_ 35nM) Clk1 (K_d_ 13nM), Clk2 (K_d_ 59nM), Clk3 (K_d_ 530nM) and CLK4 (K_d_ 26nM) (Fig. 2C; Supp. Table 2). Notably, Ribosomal protein S6 kinase A6 (RSK4), a member of the ACG family, is a false positive confirmed through subsequent full K_d’s_ determination (RSK K_d_ > 10μM, n = 2) where two full K_d_’s were sought (i.e. a dose response follow-up). Both times the K_d_ was < 10uM. RSK has not been seen in any other scan of close analogs of DYR533 further supporting it as a false positive.

**Figure 2.**
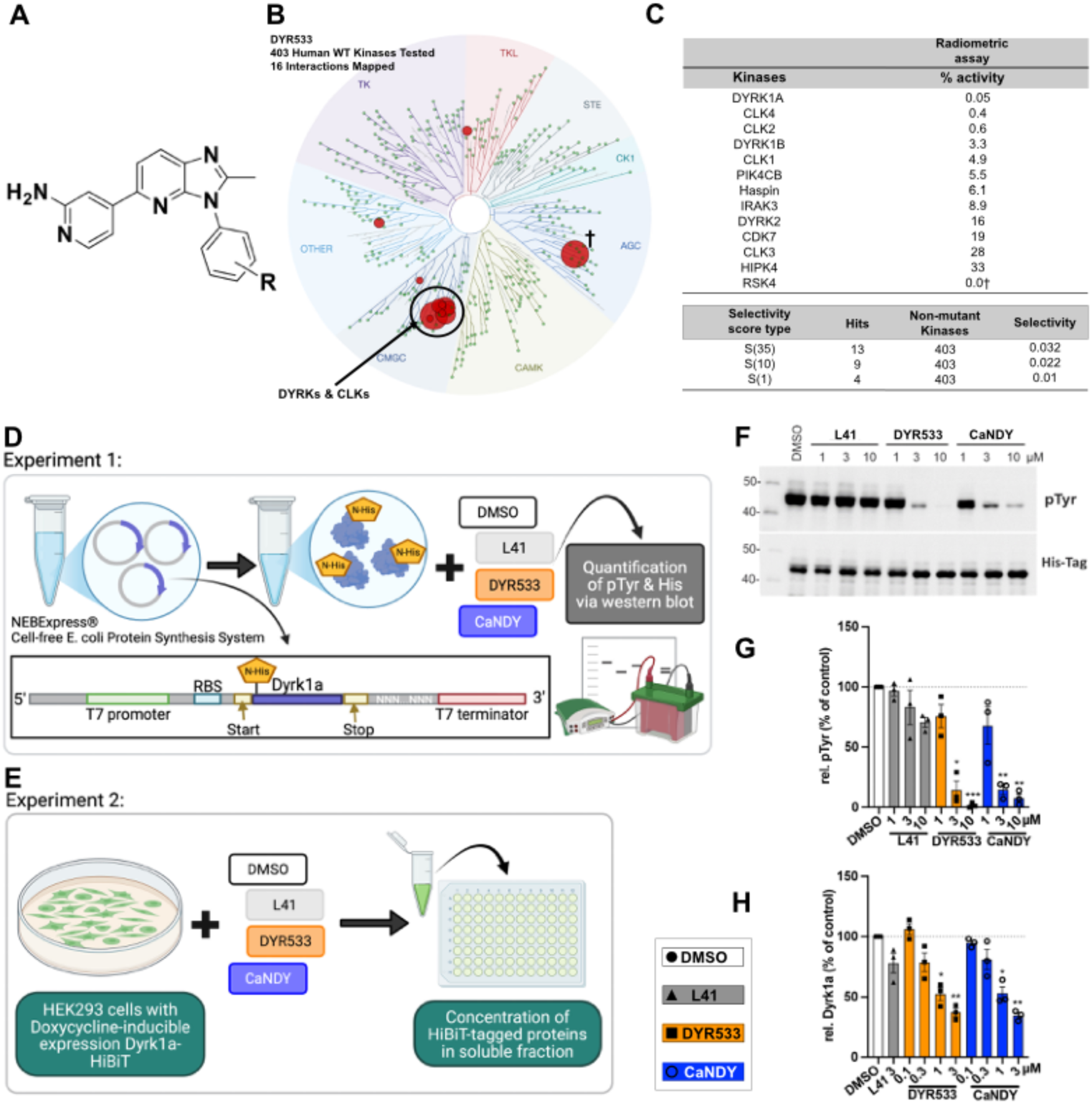
DYR533 inhibits tyrosine autophosphorylation and induces degradation of Dyrk1a. (A) Chemical structure of DYR533. Kinome profiling of DYR533 (1 μM) was conducted across 403 wildtype (WT) human kinases using the Eurofins DiscoverX KinomeScan platform (Supp. Table 2). (B) A TREEspot kinase map illustrates binding interactions, with larger circles indicating stronger binding. (C) Selectivity metrics are shown for kinases with percent control <35% ((number of non-mutant kinases with %Ctrl <35) / (number of non-mutant kinases tested)). S-scores represent the fraction of kinases bound at thresholds of <1, <10, and <35, excluding mutant variants. Kinases inhibited >85% are highlighted in bold. Cell-free and in vitro assays confirmed that DYR533 blocks Dyrk1a intramolecular tyrosine autophosphorylation—similar to the established Dyrk1a inhibitor CaNDY—and promotes Dyrk1a degradation. (D) Experimental design for phosphotyrosine (pTyr) assays and (E) Dyrk1a degradation assays. (F) Western blot results from the pTyr assay and (G) corresponding quantification showing reduced pTyr levels following treatment with DYR533 or CaNDY at 3 an 10 μM. (H) Dyrk1a degradation assay results demonstrating decreased Dyrk1a levels after treatment with 1 or 3 μM DYR533 or CaNDY. Data are presented as mean ± SEM. *p < .05, **p<0.01, p<0.0001.

Dyrk1a is autophosphorylated during the folding process at Tyrosine 321 (Tyr321)^72^, allowing the catalytic domain of the protein to acquire a mature conformation, rendering it active towards serine and threonine residues^72^. Inhibiting the autophosphorylation of Tyr321 is expected to inhibit Dyrk1a irreversibly, resulting in the degradation of the inactive protein^73^. Using a cell-free expression system (Fig. 2D), we quantified pTyr in the presence of DYR533. For comparison, we included L41 a well characterized Type 1 Dyrk1a inhibitor^39,56^, and CaNDY as an inhibitor that disrupts folding of Dyrk1a^57^. DYR533 and CaNDY significantly reduced pTyr compared to untreated control samples (Fig. 2G; *p =* 0.0049) at 3 μM (*p =* 0.0228 and *p =* 0.0062, respectively) and 10 μM (*p =* 0.0005 and *p =* 0.0044, respectively, Fig. 2F). However, no reduction was observed with L41 at 1, 3, or 10 μM (*p = 0*.9311, *p* = 0.7862, *p = 0*.0532, respectively). This suggests that DYR533 and CaNDY, but not L41, inhibit the tyrosine autophosphorylation of Dyrk1a.

To examine whether degradation of Dyrk1a occurs in the presence of DYR533, HEK293 cells stably expressing HiBiT-tagged Dyrk1a were incubated with Dyrk1a inhibitors at 0.1.-3.0 µM concentrations. The 11-amino-acid HiBiT peptide complements with LgBiT protein to produce luminescence directly proportional to the amount of the HiBiT tagged Dyrk1a protein in a split luciferase assay^74^. After 24 hours, the amount of soluble HiBiT-tagged Dyrk1a in the cell lysates was measured by a fragment complementation luciferase assay (HiBiT lytic assay; Fig. 2E). DYR533 and CaNDY reduced relative Dyrk1a protein levels (Fig. 2H; *p =* 0.0123) compared to DMSO control at 1μM (*p =* 0.0380 and *p =* 0.0411, respectively) and 3μM (*p =* 0.0093 and *p =* 0.0039, respectively). These results highlight the selectivity of DYR533 and its ability to disrupt the autophosphorylation step required for newly synthesized Dyrk1a to become active, leading to the degradation of the protein.

### DYR533 reduces Dyrk1a protein levels, AD-like pathogenesis and TNF-⍺ in the 3xTg-AD mouse model of AD

We previously tested an analog of DYR533, DYR219, in 3xTg-AD mice. Therefore, we first evaluated DYR533 in this same AD model^41,75^. We tracked weight and found a genotype effect for percent change from start of dosing (baseline) to the end of study (Fig. 3B; *p =* 0.0016), where 3xTg-AD mice lost more weight than NonTg. We found elevated Dyrk1a protein levels in Hp of 3xTg-AD Veh mice (3xVeh) compared to NonTg. These levels were reduced in a dose-dependent manner in 3xTg-AD mice administered DYR533 (3xDYR) (Fig. 3C; *p <* 0.0001). In the Ctx, a significant effect of genotype (*p* < 0.0001), dose (*p* < 0.0001), and a genotype by dose interaction (*p* < 0.0001) were found (Fig. 3D). 3xTg-AD mice showed higher Ctx Dyrk1a levels compared to NonTg, and all 3xDYR showed reduced levels compared to 3xVeh (*p* < 0.0001). Dyrk1a levels in 3xDYR 5.0 and 2.5 mg/kg were also reduced in comparison to 3xDYR 1.0 mg/kg (*p* < 0.0001), collectively highlighting efficacy of DYR533 to reduce its biological target.

**Figure 3:**
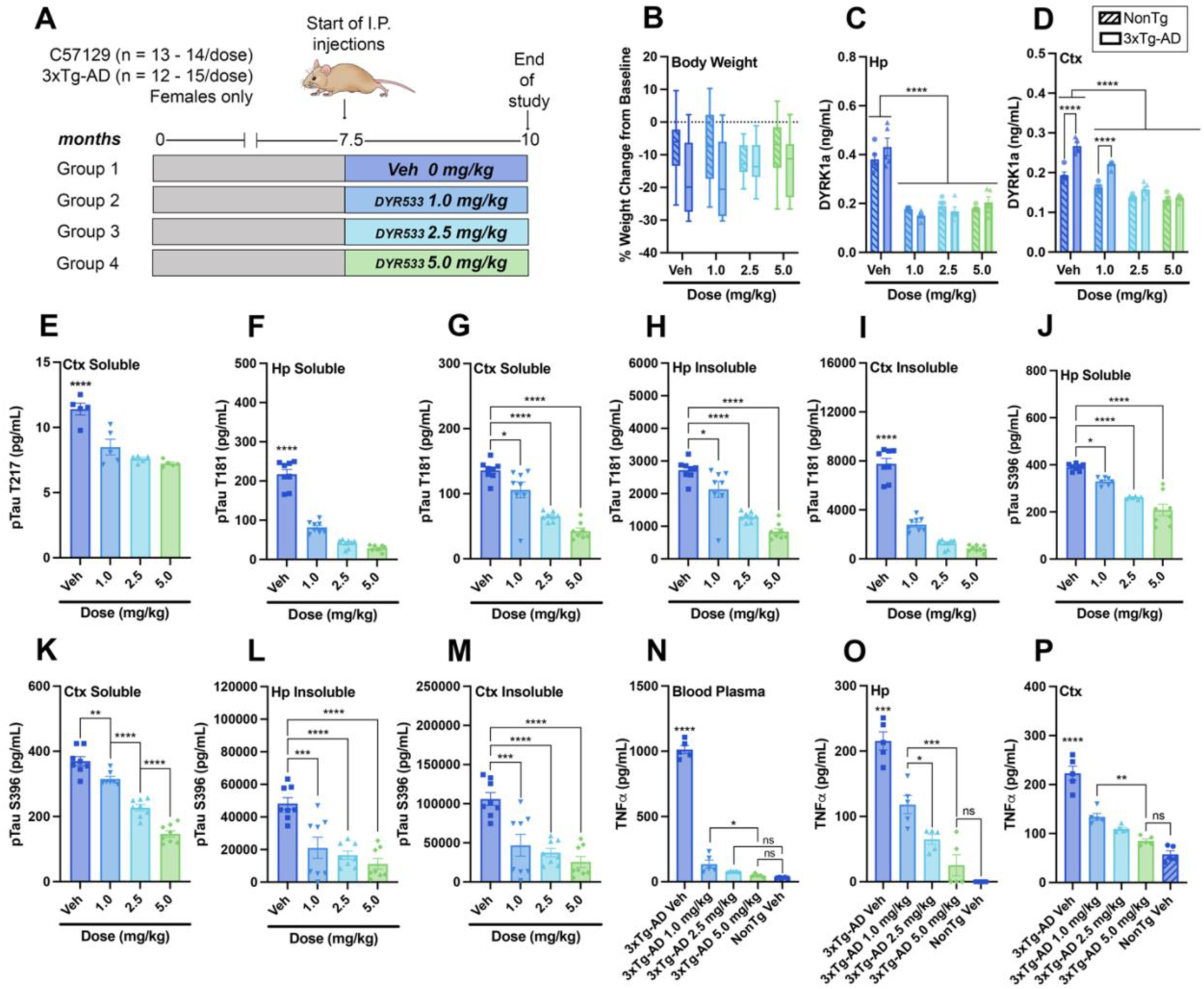
In the 3xTg-AD mouse model of AD, DYR533 reduced tau hyperphosphorylation at pathologically relevant epitopes in the hippocampus (Hp) and cortex (Ctx) and TNF-⍺ levels in the Hp, Ctx, and blood plasma. (A) Timeline of the DYR533 study in 3xTg-AD mice. (B) % weight change in groups from start of dosing (baseline) through the end of the study. (C, D) Dyrk1a protein quantifications from hippocampal (Hp) and cortical (Ctx) tissue across groups. Quantification of pTau T217 in Ctx (E), and pTau T181 (F – I) and pTau S396 (J – M) in the Hp and Ctx across 3xTg-AD groups. Quantifications of TNF-⍺ _in blood plasma (N), Hp (O), and Ctx (P) across 3xTg-AD dosed and NonTg Veh groups. For box plots, the center line represents the median value, the limits represent the 25th and 75th percentile, and the whiskers represent the minimum and maximum value of the distribution. Bar graphs are means ± SEM. \**p < 0*.05, *\*\* p* < 0.01, *\*\*\* p* < 0.001, *\*\*\*\* p* < 0.0001.

We hypothesized that DYR533 would reduce pTau levels, as Dyrk1a phosphorylates tau at pathologically relevant epitopes. Human pTau epitopes were not detectable in NonTg mice lacking human tau transgenes, thus these animals were excluded from pTau analysis, consistent with prior reports^45,47,62^. pTau at T217^76,77^ was reduced in soluble Ctx of 3xDYR mice compared to 3xVeh (Fig. 3E; *p <* 0.0001). Soluble Hp (Fig. 3F; *p <* 0.0001) pTau T181 was also reduced in 3xDYR, where 3xDYR 5.0 (*p <* 0.0001) and 2.5 (*p =* 0.0016) showed significant reductions compared to 3xDYR 1.0 mg/kg. Soluble Ctx (Fig. 3G; *p <* 0.0001) from 3xDYR showed reductions in pTau T181 compared to 3xVeh (5.0 (*p* < 0.0001), 2.5 (*p <* 0.0001), and 1.0 mg/kg (*p =* 0.0303)), and at higher doses (3xDYR5.0 (*p <* 0.0001) and 2.5 *(p* = 0.0022)) were lower than 3xDYR 1.0 mg/kg. Insoluble fractions of pTau T181 were also reduced in the Hp (Fig. 3H; *p <* 0.0001) and Ctx (Fig. 3I; *p <* 0.0001). 3xDYR had lower pTau T181 in insoluble Hp fractions compared to 3xVeh (5.0 (*p* < 0.0001), 2.5 *(p* < 0.0001), 1.0 mg/kg (*p* = 0.0335)). However, 3xDYR 1.0 mg/kg maintained higher levels compared to 5.0 (*p <* 0.0001) and 2.5 (*p =* 0.0017). All 3xDYR showed reductions in Ctx insoluble T181 compared to 3xVeh (*p <* 0.0001). Another pathologically relevant tau epitope in moderate stages of AD^78^, directly phosphorylated by Dyrk1a is S396^18,19,79^. Treatment with DYR533 reduced soluble Hp (Fig. 3J; *p <* 0.0001) and Ctx (Fig. 3K; *p <* 0.0001) pTau S396. When compared to 3xVeh, 3xDYR showed reductions in Hp S396 (5.0, (*p <* 0.0001); 2.5, (*p* < 0.0001); 1.0 mg/kg (*p* = 0.0260)). Similarly, in soluble Ctx, S396 was significantly reduced in a dose dependent manner (5.0, (*p* < 0.0001) < 2.5, (*p <* 0.0001) < 1.0 mg/kg, (*p <* 0.0001) < Veh). Insoluble pTau S396 levels in the Hp (Fig. 3L; *p <* 0.0001) and Ctx (Fig. 3M; *p <* 0.0001) were also reduced in 3xDYR compared to 3xVeh (in Hp: 5.0, (*p <* 0.0001); 2.5, (*p <* 0.0001); 1.0 mg/kg, (*p =* 0.0006); in Ctx: 5.0-, (*p <* 0.0001); 2.5, (*p <* 0.0001); 1.0 mg/kg, (*p =* 0.0006)). This demonstrates that DYR533 treatment robustly reduced phosphorylation of multiple disease-associated pTau epitopes across disease stages in soluble and insoluble Hp and Ctx fractions compared to 3xVeh.

Consistent with neuroinflammation observed in ADRD human cases, 3xTg-AD mice exhibit similar inflammatory features. Therefore, we assessed whether DYR533 treatment attenuated TNF-α expression in the brain and periphery. DYR533 reduced TNF-⍺ levels in plasma (Fig. 3N; *p <* 0.0001), Hp (Fig. 3O; *p <* 0.0001), and Ctx (Fig. 3P; *p <* 0.0001). 3xTg-AD mice also produce Aβ pathology resulting from mutated human *APP* and *PSEN1* genes. DYR533 treatment reduced Ctx soluble Aβ_42_ (Supp. Fig. 1F; *p =* 0.0007). However, reductions in Aβ40 and 42 were not significantly reduced (Supp. Fig. 1A-E, G, H). Minimal reductions in fractions of Aβ pathology may result from initiating treatment at seven and a half months of age, when Aβ is advanced. These reductions in Dyrk1a, multiple pTau epitopes, TNF-α, and soluble Aβ42 in 3xDYR versus 3xVeh demonstrate the efficacy of our novel inhibitor in a model of combined amyloid and tau pathology.

### DYR533 lowered Dyrk1a levels and markedly improved thigmotaxis and spatial memory in the PS19 tauopathy model

Given the significant reduction in tau phosphorylation observed in our 3xTg-AD study, we next tested the efficacy of DYR533 in the PS19 mouse model of tauopathy (Fig. 4A). No differences in percent weight change from baseline to end of study were found between groups (Fig. 4B). At seven and a half months of age, mice were tested on a battery of behavioral tasks. Reporting of the results from the behavioral tasks are focused on performance of all PS19 treated with DYR533 (PSDYR) and PS19 Veh (PSVeh) compared to NonTg Veh as we did not observe any negative outcomes nor benefits in NonTg mice treated with DYR533. During the two-day training period on the Rotarod task, we found a significant main effect of day (*p <* 0.0001) and group (*p =* 0.0487) for latency to fall (Fig. 4C). A trend toward significantly longer latency to fall was found between the PSDYR 1.0 mg/kg (*p =* 0.071) and 5.0mg/kg (*p =* 0.073) compared to the PSVeh. No significant differences were found in the probe day of the Rotarod (Fig. 4D). Next, animals were tested in the MWM to assess spatial learning and memory. Six animals were unable to participate due to hind-end paralysis^50^ and were excluded from testing. An additional six animals were excluded from the analysis due to high immobility (n = 5, average % immobile > 50%) or health concerns (n = 1). During the learning phase, a main effect of day (Fig. 4E; *p* < 0.0001) for latency to the platform were observed. We next calculated percent thigmotaxis, the tendency to remain near the pool edge as an indicator of anxiety^80^. We observed significant main effects of day (Fig. 4F; *p* < 0.0001) and group (*p* < 0.0001), with thigmotaxis decreasing over time. Importantly, DYR533 reduced this behavior in all PSDYR-treated mice. During the day six probe trial, we found a trend towards significance for time in platform quadrant (Fig. 4G, *p* = 0.0588), where PSVeh spent less time in the correct quadrant compared to PSDYR and NonTg Veh. Analysis of velocity did not reach significance, demonstrating that MWM performance could not be attributed to swim speed differences (Fig. 4H; *p =* 0.0714).

**Figure 4.**
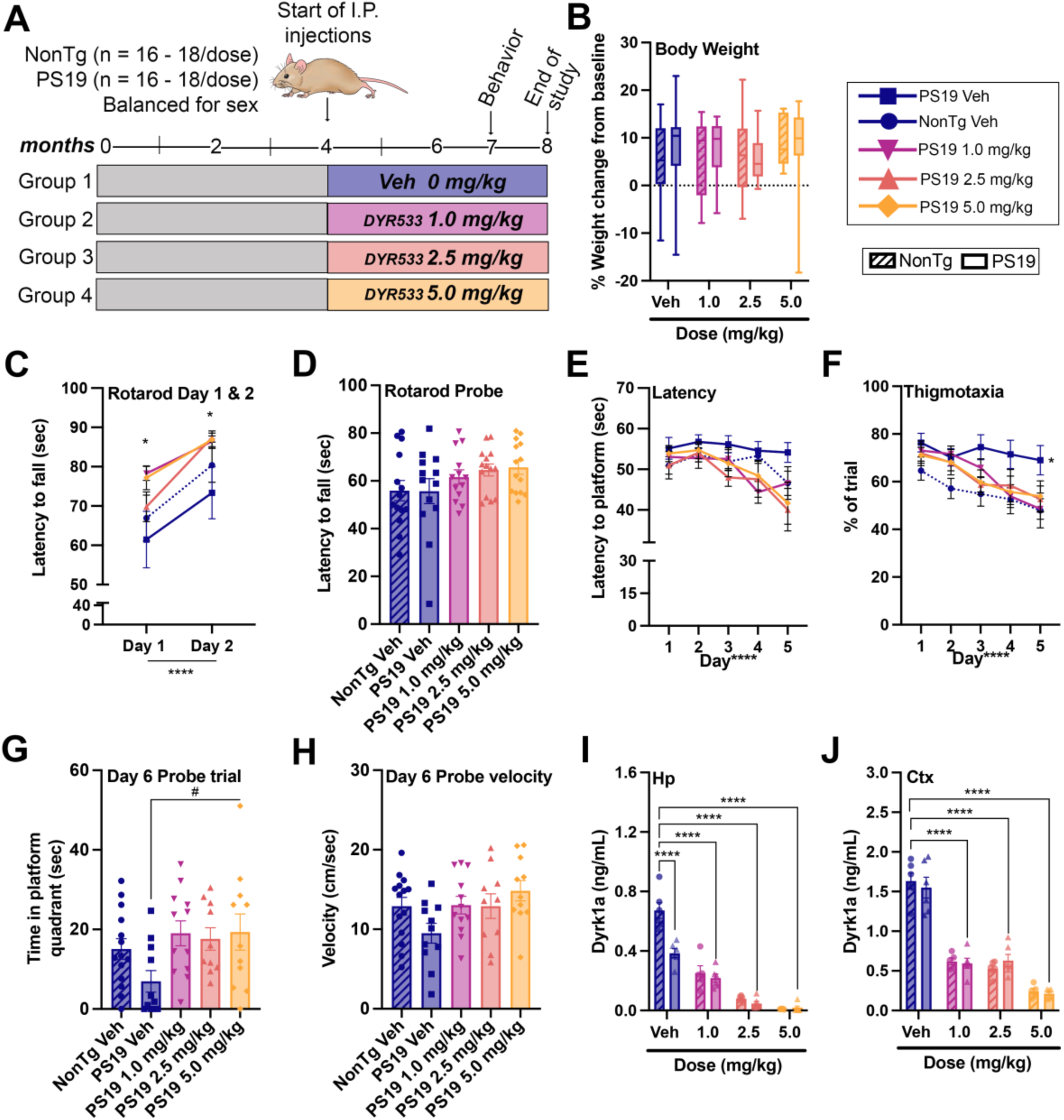
In the PS19 mouse model of tauopathy, DYR533 improved spatial learning, reduced thigmotaxia in the Morris water maze (MWM), and decreased Dyrk1a protein levels in the hippocampus (Hp) and cortex (Ctx). (A) Study timeline and experimental groups. (B) Percent weight change from baseline to study end. (C, D) Rotarod performance during training and probe sessions for PS19 groups compared with NonTg Veh. (E–H) MWM learning and memory performance across treatment groups. (I–J) Quantification of Dyrk1a protein levels from hippocampal (Hp) and cortical t (Ctx) tissue. For box plots, center lines indicate median values; boxes represent the 25th–75th percentiles; whiskers denote minimum and maximum values. Line and bar graphs depict mean ± SEM. *p < 0.05, **p < 0.01, ***p < 0.001, ****p < 0.0001, trend #p = 0.05 < 0.06.

In the Hp, we found significant main effects of genotype (*p =* 0.0003), dose (*p* < 0.0001) and genotype by dose interaction (*p* < 0.0001) (Fig. 4I) for Dyrk1a protein levels. Lower Dyrk1a levels were found in PSVeh compared to NonTg Veh mice, and DYR533 significantly lowered Dyrk1a in both PS and NonTg mice. In the Ctx, there was a significant main effect of dose (*p* < 0.0001; Fig. 4J), with DYR533 again significantly reducing Dyrk1a levels in both PS and NonTg mice. These findings demonstrate that treatment with DYR533 improved aspects of thigmotaxis in the MWM, as well as having no effect on body weight, and significantly lowering Dyrk1a levels in the Hp and Ctx of PS19 mice.

### In PS19 mice, DYR533 significantly reduced phosphorylated tau at multiple disease-associated epitopes

The P301S mutation of the *MAPT* gene is responsible for the tau pathology observed in PS19 mice, mimicking primary tauopathies. These mice produce human tau that is hyperphosphorylated at disease-relevant epitopes previously mentioned in the 3xTg-AD study. We found that Ctx soluble pTau T217 was significantly reduced (Fig. 5A; *p* < 0.0001) in all PSDYR compared to PSVeh (*p* < 0.0001). DYR533 reduced pTau at T181 in soluble Hp (Fig. 5B; *p* < 0.0001) and soluble Ctx fractions (Fig. 5C; *p* < 0.0001), highlighting that higher doses of DYR533 resulted in a greater reduction in this pTau epitope in PS19 mice (*p* < 0.05). pTau T181 was also reduced in insoluble fractions of Hp (Fig. 5D; *p* < 0.0001) and Ctx (Fig. 5E; *p* < 0.0001) in PSDYR compared to PSVeh (*p <* 0.0001). DYR533 treatment reduced Hp soluble pTau S396 (Fig. 5F; *p* < 0. 0001) in a dose dependent manner (5.0 (*p* < 0.0001) < 2.5 (*p* < 0.0001) < 1.0 mg/kg (p < 0.0001) < Veh) and Ctx soluble pTau S396 was reduced in PSDYR (Fig. 5G; *p* < 0.0001), where 5.0 (*p <* 0.0001), 2.5 (*p <* 0.0001), and 1.0 mg/kg (*p =* 0.0008) showed reduced levels compared to PSVeh. Hp insoluble pTau S396 (Fig. 5H; *p* < 0.0001) levels were also reduced in PSDYR compared to PSVeh (*p <* 0.0001). Similarly, DYR533 reduced Ctx insoluble pTau S396 (Fig. 5I; *p* < 0.0001) compared to PSVeh (5.0- (*p <* 0.0001), 2.5 (*p <* 0.0001) and 1.0 mg/kg (*p =* 0.0009)). Taken together, these data further support that DYR533 significantly reduces pTau at pathological epitopes, demonstrating its efficacy as a therapeutic for reducing tau pathogenesis in a model of primary tauopathy.

**Figure 5.**
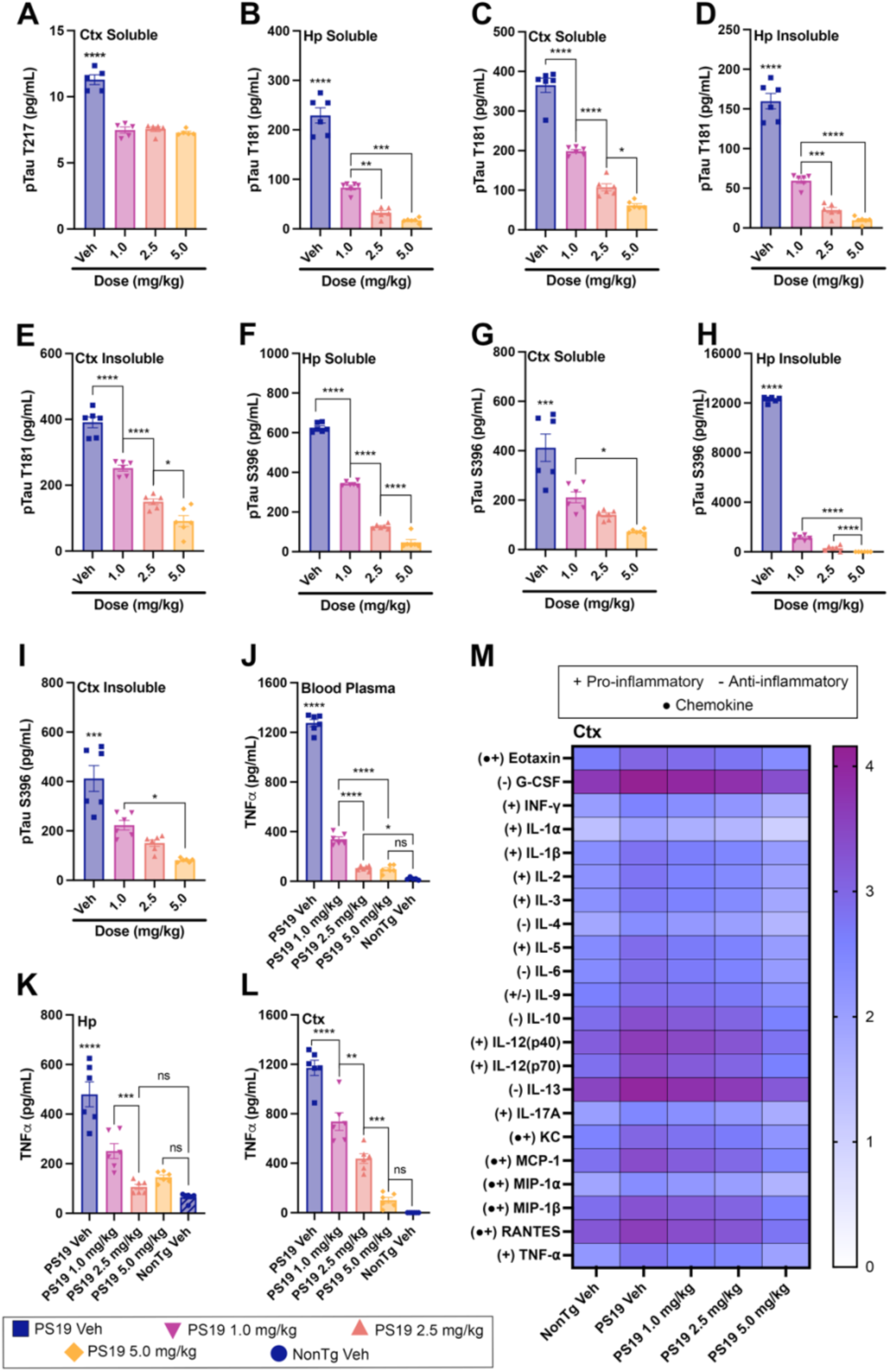
DYR533 Reduces Pathological Tau Hyperphosphorylation and Neuroinflammatory Cytokine and Chemokine Profiles in PS19 Mice. DYR533 treatment lowered tau phosphorylation at disease-relevant epitopes and reduced inflammation in blood plasma, hippocampus (Hp), and cortex (Ctx) compared with PS19 Veh. Quantification of pTau T217 in Ctx (A); pTau T181 in Hp and Ctx (B–E); and pTau S396 in Hp and Ctx (F–I). TNF-α levels in blood serum (J), Hp (K), and Ctx (L). (M) Heat map illustrating cytokine and chemokine reductions in Ctx of PS19 mice treated with DYR533, shown as log transformation from raw value. Bar graphs represent mean ± SEM. *p < 0.05, **p < 0.01, ***p < 0.001, ****p < 0.0001.

Given the associations between Dyrk1a and TNF-⍺, we next assessed the impact of DYR533 on this cytokine in PS19 mice. TNF-⍺ levels in peripheral blood plasma (Fig. 5J; *p* < 0.0001) were reduced in PSDYR compared to PSVeh. The change in TNF-⍺ levels with DYR533 treatment followed the same reductions in Hp (Fig. 5K; *p* < 0.0001) and Ctx (Fig. 5L; *p* < 0.0001), with 5.0 mg/kg showing no difference from NonTg (*p* > 0.2). These results highlight that DYR533 markedly lowered TNF-⍺ in blood plasma, Hp, and Ctx, restoring levels at the highest dose to those seen in NonTg Veh mice.

To determine whether DYR533 reduced other inflammatory molecules observed in disorders such as AD and FTD, we next used a Bio-Rad multiplex assay to quantify 23 different cytokines and chemokines simultaneously in PS19 Ctx homogenates. Out of 23 cytokines and chemokines, one cytokine, GM-CSF, was excluded because levels were outside the limit of detection. 22 of the measured inflammatory molecules were significantly elevated in PSVeh compared to NonTg Veh (*p <* 0.0001). PSDYR, at all doses, showed reductions in 18 out of the 22 cytokines and chemokines measured (Supp. Table 1 and Fig. 5M*; p <* 0.05); 5.0 and 2.5 mg/kg groups showed significant reductions in 22/22 compared to PSVeh *p <* 0.05). PSDYR mice dosed at 5.0 and 2.5 mg/kg showed equal levels compared to NonTg Veh in 21/22 cytokines/chemokines measured - PS19 5.0 mg/kg RANTES was significantly lower than NonTg Veh (*p =* 0.0119). Collectively, these findings show that DYR533 both reduces tau hyperphosphorylation and normalizes inflammatory signaling, effectively dampening brain inflammation.

### DYR533 lowers Dyrk1a, reduces AD-like pathology, and corrects immune dysregulation in the basal forebrain—a region vulnerable in Down syndrome and AD—in Ts65Dn mice

Dyrk1a is triplicated in DS and contributes to the development of AD neuropathology^34,81,82^, highlighting DYR533 as a promising therapeutic for this disorder. Given the known sex differences in this model, in this study (Fig 6A), we analyzed sex as a variable consistent with previous reports^83,84^. We first examined weight change from baseline to euthanasia finding a main effect of genotype (Fig. 6B; *p =* 0.0012), where 3n mice weighed significantly more than 2n mice. We quantified various outputs in the BF as this brain region exhibits sex-specific elevations in Aβ and early loss of cholinergic neurons in Ts65Dn mice^60,83^. For Dyrk1a protein levels, we found a significant main effect of genotype (Fig. 6C; *p <* 0.0001, 3n higher than 2n), and dose (*p <* 0.01, DYR533 lower than Veh). We also found a significant genotype by sex interaction (p = 0.047) and genotype by dose interaction (p < 0.0001). The genotype by sex post-hocs revealed that 3n females (F) (*p =* 0.0262) differed from 2n F. Genotype by dose post-hoc analyses showed that 2n Veh mice (2nVeh) (p < 0.0001) and 3n mice treated with DYR533 (3nDYR) (*p <* 0.0001) had lower Dyrk1a compared to 3n Vehicle mice (3nVeh).

**Figure 6.**
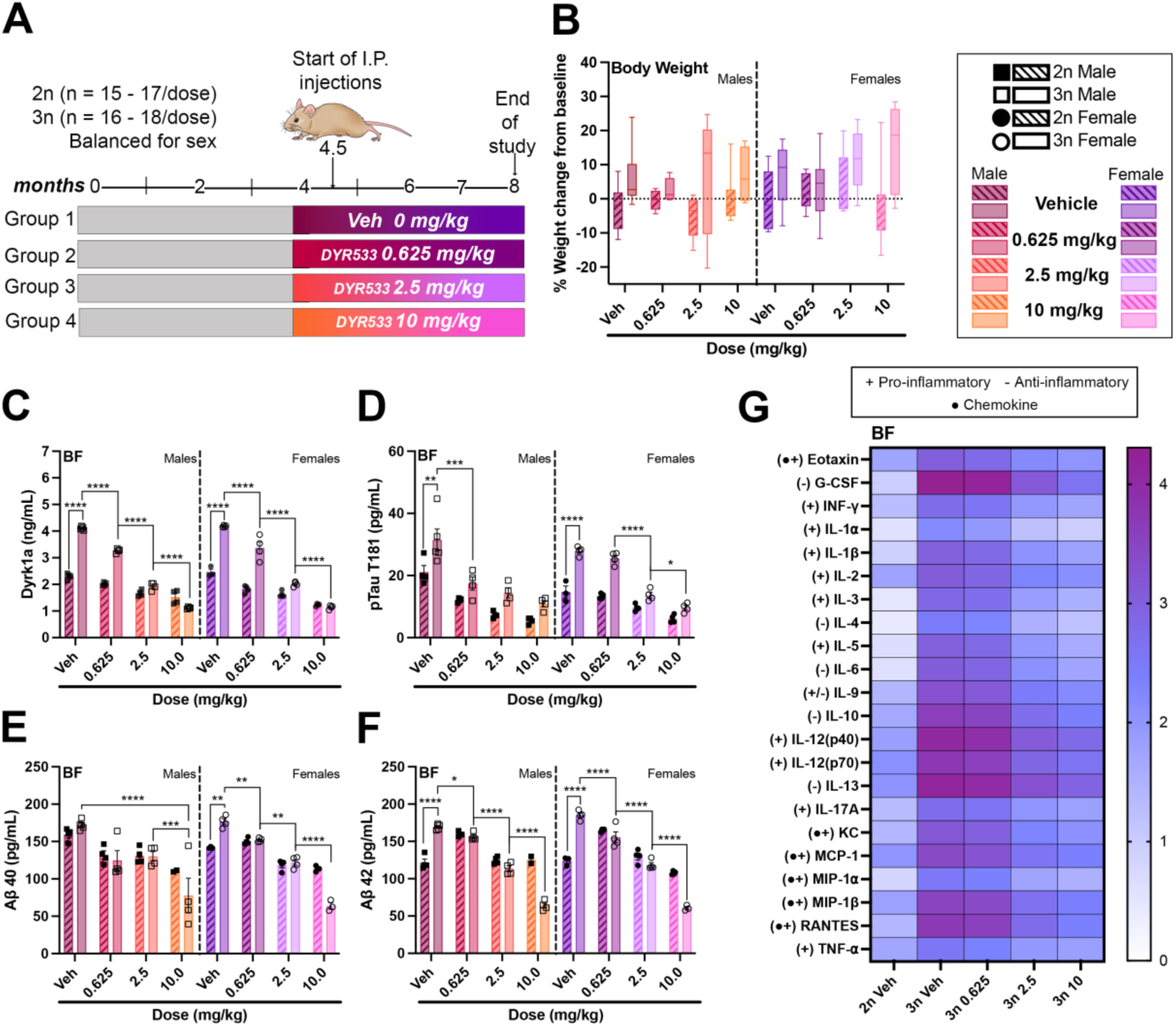
While DYR533 did not affect body weight, it reduced Dyrk1a, pTau T181, and Aβ40–42 levels in the basal forebrain of Ts65Dn mice. (A) Study timeline and groups. (B) Percent weight change from baseline to study end. Quantification of BF Dyrk1a (C), pTau T181 (D), Aβ40 (E), and Aβ42 (F). (G) Heat map showing significant reductions in 22 cytokines and chemokines in the BF of 3n mice treated with DYR533, expressed as log transformation from raw value. For box plots, the center line denotes the median, box limits represent the 25th and 75th percentiles, and whiskers indicate the minimum and maximum values. Bar graphs represent mean ± SEM. *p < 0.05, **p < 0.01, ***p < 0.001, ****p < 0.0001.

While Ts65Dn do not develop Aβ and tau aggregates, quantification of soluble murine pTau T181 and Aβ_40_ and Aβ_42_ from BF homogenates is possible. We found a main effect of genotype (Fig. 6D; *p <* 0.0001, 3n higher than 2n), dose (*p <* 0.0001, DYR533 lower than Veh), and genotype by dose (*p =* 0.0092) interaction for pTau T181. 2nVeh (*p <* 0.0001) and 3nDYR (0.625 (*p =* 0.0003), 2.5 (*p <* 0.0001), and 10 mg/kg (*p <* 0.0001)) had significantly lower pTau T181 levels compared to 3nVeh. While a significant sex by dose (*p =* 0.0013) interaction was found, post-hoc comparisons did not reach statistical significance. Next, we quantified Aβ_40_ levels and found a main effect of dose (Fig. 6E; *p <* 0.0001), where all DYR treated mice showed significantly lower levels, and the 10mg/kg group showed lower levels than the 0.625mg/kg (*p <* 0.0001) and 2.5mg/kg (*p <* 0.0001) groups. We also found a genotype by dose (*p <* 0.0001) interaction, where 3n 0.625mg/kg and 2.5mg/kg levels were higher than 2n 0.625mg/kg (*p <* 0.0001) and 2.5mg/kg (*p =* 0.0234). Notably, 2n 10mg/kg did not differ from 3n 10mg/kg. A sex by dose (*p =* 0.0156) interaction was also found, however post-hoc comparisons failed to reach statistical significance. We also quantified Aβ_42_ levels in the BF and found a main effect of genotype (Fig. 6F; *p =* 0.0182, 3n higher than 2n) and dose (*p <* 0.0001, DYR533 lower than Veh), where 2.5mg/kg (*p <* 0.0001) and 10mg/kg (*p <* 0.0001) where lower than both the Veh and 0.625mg/kg doses, and 10mg/kg was lower than 2.5mg/kg (*p =* 0.0001). A significant genotype by dose interaction (*p <* 0.0001) revealed that 3n 2.5 mg/kg differed from 2n 2.5 mg/kg (*p =* 0.0216), and 3n 10 mg/kg differed from 2n 10 mg/kg (*p <* 0.0001). Finally, a significant sex by dose interaction was found (*p =* 0.0057), but post-hoc comparisons failed to reach statistical significance. These results highlight the ability of DYR533 to lower soluble levels of Aβ in the BF of Ts65Dn mice.

Lastly, we examined the effects of DYR533 on cytokine and chemokine levels in the BF (Fig. 6G). Out of the 23 cytokines and chemokines tested, we detected 22, as GM-CSF levels were outside the limit of detection. We found a significant main effect of group (Supp. Table 1), but no significant effect of sex or interaction. Individual graphs in Supp. Fig. 3 highlight that 3n 2.5 and 10 mg/kg groups did not significantly differ in 22/22 cytokines and chemokines, suggesting that treatment at the highest dose (10mg/kg) of DYR533 is not necessary to rescue levels of cytokines and chemokines. Collectively, these data demonstrate that DYR533 significantly reduces neuroinflammation in addition to reducing tau hyperphosphorylation and Aβ_40-42_ pathology in the vulnerable BF of Ts65Dn mice.

## Discussion

We observed increased Dyk1a protein levels in primary tauopathies—including Pick’s disease, consistent with previously work^11^, and in CBD, and PSP—and these elevations were inversely correlated with MMSE scores and positively correlated with Braak stage. Consistent with previous findings^11,12^, we also found that Dyrk1a protein expression levels were elevated in human AD brains in a disease-associated manner. Notably, we found significant elevations of Dyrk1a in blood serum from AD Sev compared to age-matched control cases, but not in AD Mod, highlighting that serum Dyrk1a levels may not be a reliable biomarker across disease stages, especially in earlier stages. We further observed positive correlations between TNF-α and Dyrk1a levels in both blood and brain tissue. Using mouse models of AD, tauopathy, and DS, we demonstrated the therapeutic potential of DYR533 in mitigating tau pathology, neuroinflammation, amyloidosis, thigmotaxis and spatial cognition. These results highlight Dyrk1a dysregulation in ADRD and support the use of Dyrk1a inhibitors, like DYR533, as therapeutic interventions for neurodegenerative diseases.

DYR533 prevented the autophosphorylation of tyrosine, similarly to CaNDY, but not L41, effectively rendering the kinase inactive. A common limitation of kinase inhibitors is their promiscuity^85^, making the high specificity of DYR533 (S(35) selectivity score of 0.027 at 1μm) particularly desirable compared to CaNDY. The next closest kinases inhibited by DYR533 are Dyrk1b, CLK1, and CLK4, but at a 10x lower affinity. CaNDY inhibits Dyrk1b, CLK1, CLK2, and Haspin, which phosphorylates histone H3, by over 90% at 1μm^57^. Additionally, because DYR533 is an ATP-competitive inhibitor, it also blocks the ATP-binding site of Dyrk1a preventing kinase function. We demonstrate that DYR533 promotes Dyrk1a degradation *in vitro* and observed significant reductions in Dyrk1a levels in 3xTg-AD, PS19, and Ts65Dn mice treated with the drug. Dyrk1a has previously been shown to be degraded through the E3 ubiquitin ligase SCF^βTrCP^ in HEK293 cells^86^. In 2019, Velazquez, et al., showed that DYR219, a compound similar to DYR533, reduced Dyrk1a protein via the proteosome. Our present findings suggest that DYR533 may act through a shared cellular protein-disposal mechanism across disease models.

Both 3xTg-AD and Ts65Dn mice treated with DYR533 showed reductions in Aβ. Previous work has shown that Dyrk1a overexpression increases the affinity of APP to BACE1, through phosphorylation of APP and Presenilin 1 (PSEN1), leading to increased β-secretase cleavage^12,87^. In the 3xTg-AD study, dosing was initiated after Aβ pathology was already established. Thus, while DYR533 treatment at this later disease stage may help slow further accumulation of Aβ plaques by reducing APP phosphorylation, it is unlikely to enhance clearance of existing Aβ monomers or oligomers. Conversely, in our Ts65Dn study, dosing began at four and a half months of age, when Aβ pathology is still in its early stages^60^ - and we found greater reductions in Aβ_40_ and Aβ_42_ in the BF. Early treatment with DYR533 likely minimizes Aβ pathology by reducing APP and PSEN1 phosphorylation, which may in turn delay the onset of tau pathology^41,88–90^. Importantly, the 3xTg-AD study emphasizes that, even when Aβ pathology has progressed, DYR533 significantly reduces tau hyperphosphorylation.

Tau hyperphosphorylation is a hallmark of various neurodegenerative diseases, yet there are currently no effective therapies that directly target aberrant tau. We demonstrate that DYR533 treatment reduced the phosphorylation of tau at T181 and S396^18,19^, sites that are pathological epitopes in AD, tauopathies, and DS. However, there could be alternative ways in which DYR533 mitigated tau pathology. Previous studies have shown that Dyrk1a overexpression increases tau mRNA stability, thereby reducing its degradation and leading to elevated tau expression^13^. Tau exists in six isoforms, that vary by the number of N-terminal inserts and by the presence of either three or four microtubule-binding repeats (3R or 4R), determined by inclusion or exclusion of exon 10^91^. An imbalance in 3R:4R tau isoform expression is a characteristic feature of AD, FTD, and DS^91,92^. Dyrk1a overexpression promotes exon 10 exclusion^93,94^, thereby increasing 3R tau levels. Therefore, inhibiting Dyrk1a may help preserve exon 10 inclusion and mitigate tau isoform imbalance driving pathogenesis. Additionally, Dyrk1a primes other kinases, including GSK-3β, which phosphorylates tau^95^. DYR533 may also reduce pTau levels indirectly by modulating GSK-3β activity. Together, the combined effects of Dyrk1a inhibition-reducing tau phosphorylation, preventing exon 10 exclusion, and decreased priming of kinases such as GSK-3β, offer a plausible explanation for the robust reduction in pathological tau species observed in these studies.

Another possibility is that DYR533 potentially altered autophagy, the cellular process responsible for clearing of aberrant proteins and cellular debris^96^, which is necessary for the degradation of pathological tau^97–99^. Previous work has shown that Dyrk1a overexpression increases the phosphorylation of Tuberous sclerosis complex 2 (TSC2), hyperactivating mammalian target of rapamycin (mTOR), a mediator of autophagy, and preventing the initiation of this process^100^. DYR533 may reduce the phosphorylation of TSC2, improving mTOR-regulated autophagy and increasing clearance of pTau. Another plausible explanation is that multiple mechanisms act together: reduced pTau, preserved exon 10 splicing, decreased hyperphosphorylation by GSK-3β, and improved autophagy may collectively contribute to the attenuation of tau pathogenesis. Future work will examine the effects of DYR533 on autophagy to separate which mechanism(s) are driving the reduction in tau observed across these studies.

An interesting and unexpected result of DYR533 treatment across all models, and another possible mechanism mitigating tau hyperphosphorylation, was the significant reduction of systemic inflammatory molecules. Although Dyrk1a is highly expressed in microglia^10^, its specific role in this cell type remains poorly understood. Some studies have indicated that Dyrk1a plays a role in peripheral inflammatory signaling^101,102^ and contributes to immune responses in mouse models of LPS-induced neuroinflammation^22,23^. Across all models, we observed significant reductions in TNF-α levels in both blood plasma and brain tissue. These findings support the potential use of circulating blood TNF-α for monitoring therapeutic response to DYR533 treatment. Moreover, in cortical homogenates from the PS19 and BF homogenates from Ts65Dn mice, we found a significant reduction in multiple pro- and anti-inflammatory cytokines and chemokines. Neuroinflammation typically precedes the development of AD^103^ and exacerbates tau pathology in AD and other tauopathies^104–108^. For example, IL-1β activates kinases such as p38-mitogen activating protein kinase (MAPK) and GSK-3β^107^, which both promote tau phosphorylation and aggregation^109^. Blocking IL-1 signaling, via IL-1 receptor antagonists, can improve cognition^110^ and attenuate tau pathology in mouse models of ADRD by reducing the activity of IL-1β-dependent tau kinases, including—GSK-3β, which is primed by Dyrk1a, and CDK-5^106^. Additionally, IL-17 disrupts BBB, facilitating infiltration of T helper 17 (Th17) cells, which can damage neurons and further increase expression of pro-inflammatory IL-17, −21, and −22^111^. Conversely, treatment of Ts65Dn mice with an anti-IL17 antibody partially improved cognitive function, reduced cytokine expression and microglia density, and normalized Aβ levels^112^. The robust anti-inflammatory effect observed with DYR533 indicates the need to further delineate the role of Dyrk1a in neuroinflammation, as it remains unresolved whether these reductions stem from decreased pathological burden or from direct regulation of microglial function.

Our findings reveal that DYR533 may hold therapeutic potential for ADRD, where Dyrk1a levels are elevated in the brain. In our studies, DYR533 treatment was associated with reductions in tau pathology, amyloidosis, and neuroinflammation. While many previously developed Dyrk1a inhibitors have lacked selectivity, potentially contributing to off-target effects and limiting their translational potential, DYR533 demonstrated greater potency and selectivity in our preclinical assessments. These findings suggest that modulation of Dyrk1a can influence multiple disease-relevant processes, although further work is necessary to elucidate the entirety of underlying mechanisms and to determine their relevance in human disease.

## Supporting information

Supplemental Table 1 Statistical Outputs

Supplemental Table 2 Kiome Scan Raw data

## Acknowledgements

We would like to thank Silvia Detro-Dassen for their technical assistance and Ulli Rothweiler (Arctic University of Norway, Tromsø) for kindly providing pEXP17-DYRK1A. Figure 2 was partly created by BioRender.

## Data Sharing

Data will be made available upon reasonable request.

## Author contributions

S.K.B. Experimentation, data analysis, wrote the manuscript. J.T. Experimentation, wrote and edited the manuscript. W.W. Elisa, data analysis, edited the manuscript. C.F. Development of drug, data analysis, edited manuscript. A.S. Development of drug, data analysis, edited manuscript. S.R. Synthesis of DYR533, edited manuscript. S.T. Animal experiments, edited manuscrtupt. J.M.J. Animal experiments, edited manuscrtupt. T.G.B. Funding, human tissue collection, edited the manuscript. G.E.S. Human tissue collection, edited the manuscript. G.W. Cell assay experiments, edited the manuscript. R.K. Cell assay experiments, edited the manuscript. A.M. Kiome scan analysis, edited the manuscript. S.G. Cell assay experiments, edited the manuscript. W.B. Funding, supervised, data analysis, wrote the manuscript. C.H. Funding, supervised, data analysis, wrote the manuscript. T.D. Funding, supervised, data analysis, wrote the manuscript. R.V. Funding, supervised, data analysis, wrote the manuscript.

## Supplementary Figures

**Supplementary Figure 1:**
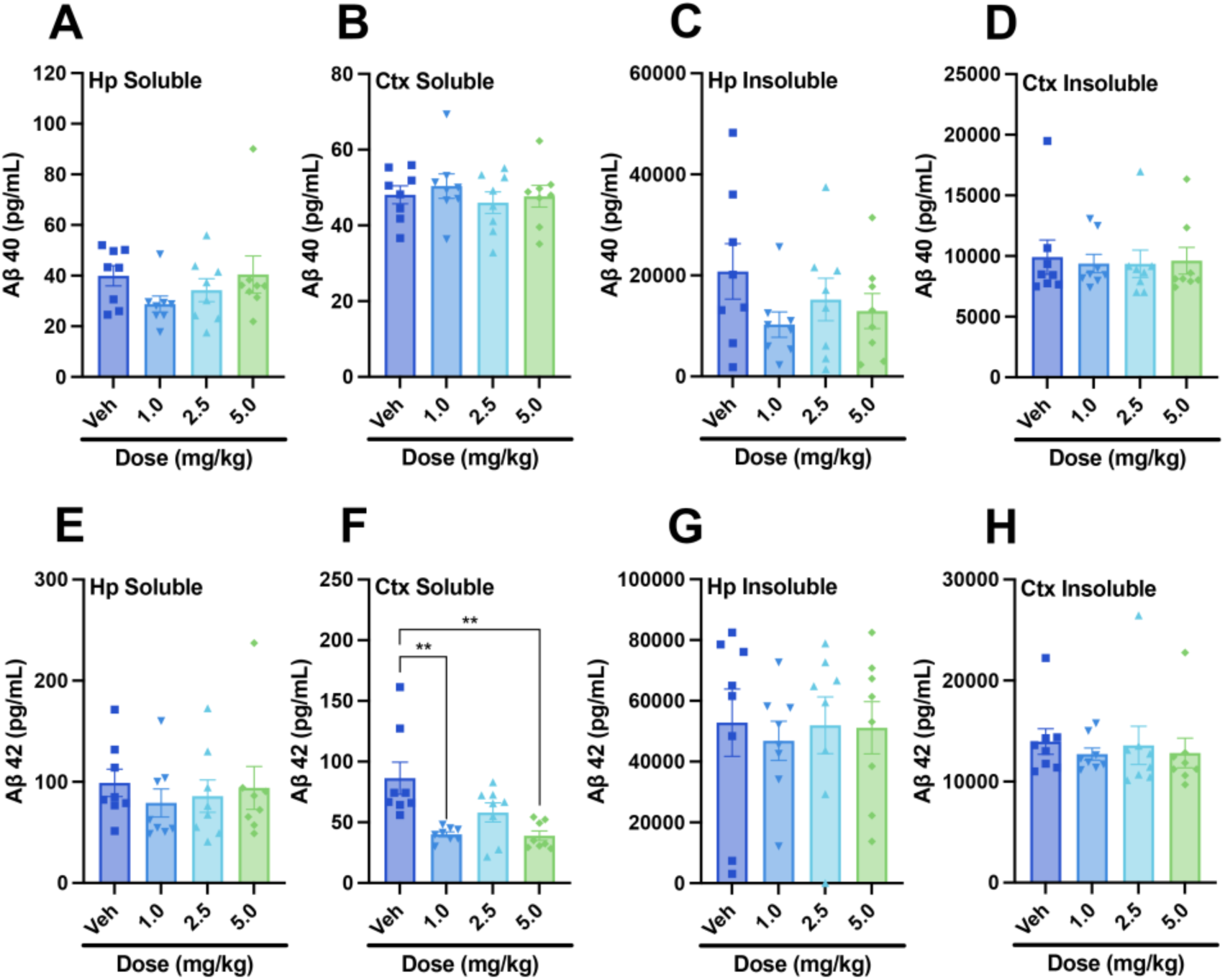
In the 3xTg-AD model, soluble Aβ42 levels in the cortex showed modest reductions, though significance was not reached for other fractions. Panels show quantifications of Aβ40 in (A) Hp soluble, (B) Ctx soluble, (C), Hp insoluble, and (D) Ctx insoluble fractions, and Aβ42 in (E) Hp soluble, (F) Ctx soluble, (G) Hp insoluble, and (H) Ctx insoluble fractions. Data are presented as mean ± SEM. *p < 0.05, **p < 0.01, ***p < 0.001, ****p < 0.0001.

**Supplementary Figure 2:**
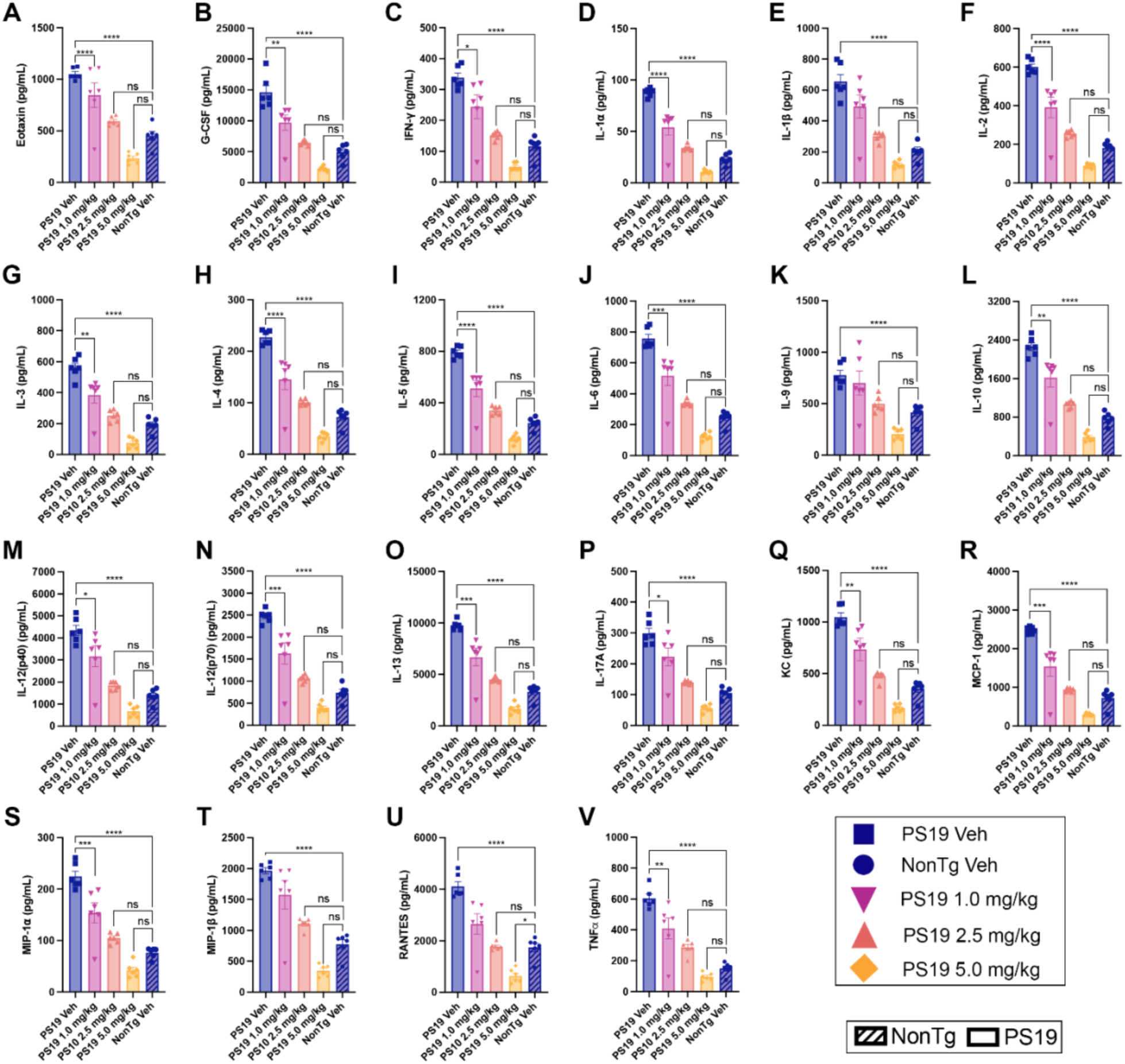
(A-V) Individual graphs from PS19 study depicting the 22 cytokines and chemokines shown in the Fig. 5M heat map and Table 1. Data are presented as mean ± SEM. *p < 0.05, **p < 0.01, ***p < 0.001, ****p < 0.0001, ns = non-significant.

**Supplementary Figure 3:**
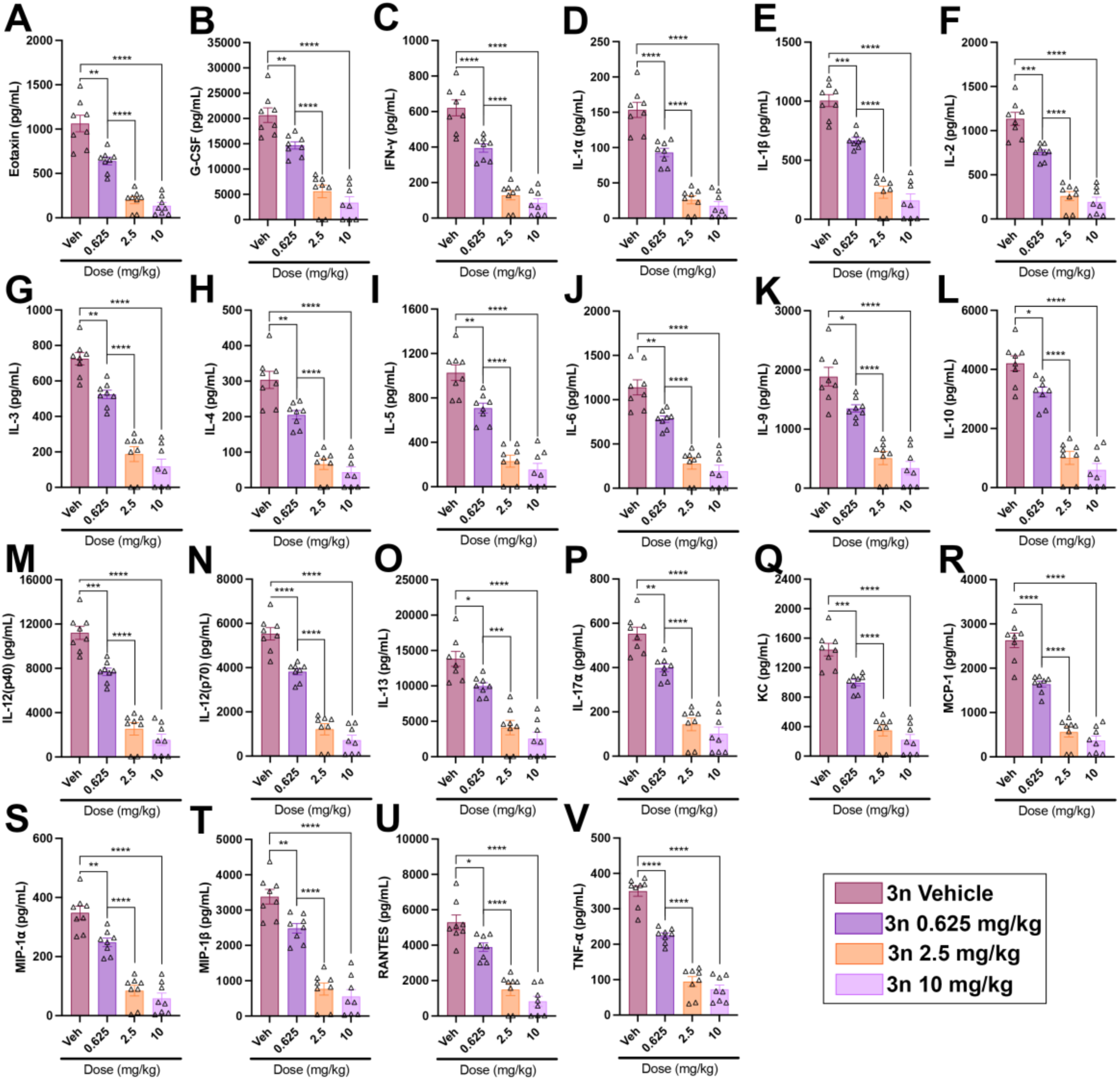
(A-V) Individual graphs from Ts65Dn study depicting the 22 cytokines and chemokines shown in the Fig. 6G heat map and Table 2. Data are presented as mean ± SEM. *p < 0.05, **p < 0.01, ***p < 0.001, ****p < 0.0001.

## References

1. Granholm, A.-C. & Hamlett, E. D. The Role of Tau Pathology in Alzheimer’s Disease and Down Syndrome. J Clin Med 13, 1338 (2024).

2. Kovacs, G. G., Ghetti, B. & Goedert, M. Classification of diseases with accumulation of Tau protein. Neuropathol Appl Neurobiol 48, e12792 (2022).

3. Perl, D. P. Neuropathology of Alzheimer’s disease. Mt Sinai J Med 77, 32–42 (2010).

4. Burger, P. C. & Vogel, F. S. The development of the pathologic changes of Alzheimer’s disease and senile dementia in patients with Down’s syndrome. Am J Pathol 73, 457–476 (1973).

5. Glenner, G. G. & Wong, C. W. Alzheimer’s disease and Down’s syndrome: Sharing of a unique cerebrovascular amyloid fibril protein. Biochemical and Biophysical Research Communications 122, 1131–1135 (1984).

6. Tanzi, R. E. et al. Amyloid beta protein gene: cDNA, mRNA distribution, and genetic linkage near the Alzheimer locus. Science 235, 880–884 (1987).

7. Aisen, P. S. et al. AHEAD 3-45 study design: A global study to evaluate the efficacy and safety of treatment with BAN2401 for 216 weeks in preclinical Alzheimer’s disease with intermediate amyloid (A3 trial) and elevated amyloid (A45 trial). Alzheimer’s & Dementia 16, e044511 (2020).

8. Hartnell, lain. Three promising drugs for treating Alzheimer’s disease bring fresh hope | Alzheimer’s Society. Alzheimer’s Society https://www.alzheimers.org.uk/blog/three-promising-drugs-for-treating-alzheimers-disease-bring-fresh-hope (2024).

9. Kabir, Md. T., et al. Combination Drug Therapy for the Management of Alzheimer’s Disease. Int J Mol Sci 21, 3272 (2020).

10. Sjöstedt, E. et al. An atlas of the protein-coding genes in the human, pig, and mouse brain. Science 367, eaay5947 (2020).

11. Ferrer, I. et al. Constitutive Dyrk1A is abnormally expressed in Alzheimer disease, Down syndrome, Pick disease, and related transgenic models. Neurobiol Dis 20, 392–400 (2005).

12. Kimura, R. et al. The DYRK1A gene, encoded in chromosome 21 Down syndrome critical region, bridges between beta-amyloid production and tau phosphorylation in Alzheimer disease. Hum Mol Genet 16, 15–23 (2007).

13. Qian, W. et al. Dual-specificity tyrosine phosphorylation-regulated kinase 1A (Dyrk1A) enhances tau expression. J Alzheimers Dis 37, 529–538 (2013).

14. Wegiel, J., Gong, C.-X. & Hwang, Y.-W. The role of DYRK1A in neurodegenerative diseases. The FEBS Journal 278, 236–245 (2011).

15. Park, J., Song, W.-J. & Chung, K. C. Function and regulation of Dyrk1A: towards understanding Down syndrome. Cell Mol Life Sci 66, 3235–3240 (2009).

16. Ryoo, S.-R. et al. Dual-specificity tyrosine(Y)-phosphorylation regulated kinase 1A-mediated phosphorylation of amyloid precursor protein: evidence for a functional link between Down syndrome and Alzheimer’s disease. J Neurochem 104, 1333–1344 (2008).

17. Ulku, I. et al. Inhibition of BACE1 affected both its Aβ producing and degrading activities and increased Aβ42 and Aβ40 levels at high-level BACE1 expression. J Biol Chem 300, 107510 (2024).

18. Bramblett, G. T. et al. Abnormal tau phosphorylation at Ser396 in Alzheimer’s disease recapitulates development and contributes to reduced microtubule binding. Neuron 10, 1089–1099 (1993).

19. Liu, F. et al. Overexpression of Dyrk1A contributes to neurofibrillary degeneration in Down syndrome. FASEB J 22, 3224–3233 (2008).

20. Ryoo, S.-R. et al. DYRK1A-mediated Hyperphosphorylation of Tau. Journal of Biological Chemistry 282, 34850–34857 (2007).

21. Song, W.-J. et al. Phosphorylation and Inactivation of Glycogen Synthase Kinase 3β (GSK3β) by Dual-specificity Tyrosine Phosphorylation-regulated Kinase 1A (Dyrk1A)*. Journal of Biological Chemistry 290, 2321–2333 (2015).

22. Ju, C. et al. Inhibition of Dyrk1A Attenuates LPS-Induced Neuroinflammation via the TLR4/NF-κB P65 Signaling Pathway. Inflammation 45, 2375–2387 (2022).

23. Latour, A. et al. LPS-Induced Inflammation Abolishes the Effect of DYRK1A on IkB Stability in the Brain of Mice. Mol Neurobiol 56, 963–975 (2019).

24. van Bon, B. et al. Intragenic deletion in DYRK1A leads to mental retardation and primary microcephaly. Clinical Genetics 79, 296–299 (2011).

25. Manubens-Gil, L. et al. Deficits in neuronal architecture but not over-inhibition are main determinants of reduced neuronal network activity in a mouse model of overexpression of Dyrk1A. Cereb Cortex 34, bhad431 (2024).

26. Brault, V. et al. Dyrk1a gene dosage in glutamatergic neurons has key effects in cognitive deficits observed in mouse models of MRD7 and Down syndrome. PLoS Genet 17, e1009777 (2021).

27. De Toma, I., Ortega, M., Aloy, P., Sabidó, E. & Dierssen, M. DYRK1A Overexpression Alters Cognition and Neural-Related Proteomic Pathways in the Hippocampus That Are Rescued by Green Tea Extract and/or Environmental Enrichment. Front. Mol. Neurosci. 12, (2019).

28. García-Cerro, S. et al. Overexpression of Dyrk1A is implicated in several cognitive, electrophysiological and neuromorphological alterations found in a mouse model of Down syndrome. PLoS One 9, e106572 (2014).

29. Nguyen, T. L. et al. Correction of cognitive deficits in mouse models of Down syndrome by a pharmacological inhibitor of DYRK1A. Disease Models & Mechanisms 11, dmm035634 (2018).

30. Souchet, B. et al. Inhibition of DYRK1A proteolysis modifies its kinase specificity and rescues Alzheimer phenotype in APP/PS1 mice. acta neuropathol commun 7, 46 (2019).

31. Stensen, W. et al. Novel DYRK1A Inhibitor Rescues Learning and Memory Deficits in a Mouse Model of Down Syndrome. Pharmaceuticals (Basel) 14, 1170 (2021).

32. Zhu, B. et al. DYRK1A antagonists rescue degeneration and behavioural deficits of in vivo models based on amyloid-β, Tau and DYRK1A neurotoxicity. Sci Rep 12, 15847 (2022).

33. Gehlot, P., Pathak, R., Kumar, S., Choudhary, N. K. & Vyas, V. K. A review on synthetic inhibitors of dual-specific tyrosine phosphorylation-regulated kinase 1A (DYRK1A) for the treatment of Alzheimer’s disease (AD). Bioorganic & Medicinal Chemistry 113, 117925 (2024).

34. Neumann, F. et al. DYRK1A inhibition and cognitive rescue in a Down syndrome mouse model are induced by new fluoro-DANDY derivatives. Sci Rep 8, 2859 (2018).

35. Smith, B., Medda, F., Gokhale, V., Dunckley, T. & Hulme, C. Recent advances in the design, synthesis, and biological evaluation of selective DYRK1A inhibitors: a new avenue for a disease modifying treatment of Alzheimer’s? ACS Chem Neurosci 3, 857–872 (2012).

36. Hsu, M.-H. et al. Leucettamine B analogs and their carborane derivative as potential anti-cancer agents: Design, synthesis, and biological evaluation. Bioorg Chem 98, 103729 (2020).

37. Hanks, S. K. & Hunter, T. The eukaryotic protein kinase superfamily: kinase (catalytic) domain structure and classification. The FASEB Journal 9, 576–596 (1995).

38. Henderson, S. H. et al. Mining Public Domain Data to Develop Selective DYRK1A Inhibitors. ACS Med. Chem. Lett. 11, 1620–1626 (2020).

39. Tahtouh, T. et al. Selectivity, cocrystal structures, and neuroprotective properties of leucettines, a family of protein kinase inhibitors derived from the marine sponge alkaloid leucettamine B. J Med Chem 55, 9312–9330 (2012).

40. Jarhad, D. B., Mashelkar, K. K., Kim, H.-R., Noh, M. & Jeong, L. S. Dual-Specificity Tyrosine Phosphorylation-Regulated Kinase 1A (DYRK1A) Inhibitors as Potential Therapeutics. J. Med. Chem. 61, 9791–9810 (2018).

41. Velazquez, R. et al. Chronic Dyrk1 Inhibition Delays the Onset of AD-Like Pathology in 3xTg-AD Mice. Mol Neurobiol 56, 8364–8375 (2019).

42. Beach, T. G. et al. Arizona Study of Aging and Neurodegenerative Disorders and Brain and Body Donation Program. Neuropathology 35, 354–389 (2015).

43. Judd, J. M. et al. Inflammation and the pathological progression of Alzheimer’s disease are associated with low circulating choline levels. Acta Neuropathol 146, 565–583 (2023).

44. Consensus Recommendations for the Postmortem Diagnosis of Alzheimer’s Disease. Neurobiology of Aging 18, S1–S2 (1997).

45. Velazquez, R. et al. Lifelong choline supplementation ameliorates Alzheimer’s disease pathology and associated cognitive deficits by attenuating microglia activation. Aging Cell 18, e13037 (2019).

46. Winslow, W. et al. IntelliCage Automated Behavioral Phenotyping Reveals Behavior Deficits in the 3xTg-AD Mouse Model of Alzheimer’s Disease Associated With Brain Weight. Front Aging Neurosci 13, 720214 (2021).

47. Bartholomew, S. K. et al. Glyphosate exposure exacerbates neuroinflammation and Alzheimer’s disease-like pathology despite a 6-month recovery period in mice. Journal of Neuroinflammation 21, 316 (2024).

48. Dennison, J. L., Ricciardi, N. R., Lohse, I., Volmar, C.-H. & Wahlestedt, C. Sexual Dimorphism in the 3xTg-AD Mouse Model and Its Impact on Pre-Clinical Research. J Alzheimers Dis 80, 41–52 (2021).

49. Velazquez, R. et al. Central insulin dysregulation and energy dyshomeostasis in two mouse models of Alzheimer’s disease. Neurobiol Aging 58, 1–13 (2017).

50. Yoshiyama, Y. et al. Synapse Loss and Microglial Activation Precede Tangles in a P301S Tauopathy Mouse Model (DOI:10.1016/j.neuron.2007.01.010). Neuron 54, 343–344 (2007).

51. Ahmed, M. M., Sturgeon, X., Ellison, M., Davisson, M. T. & Gardiner, K. J. Loss of correlations among proteins in brains of the Ts65Dn mouse model of down syndrome. J Proteome Res 11, 1251–1263 (2012).

52. Antonarakis, S. E., Lyle, R., Dermitzakis, E. T., Reymond, A. & Deutsch, S. Chromosome 21 and down syndrome: from genomics to pathophysiology. Nat Rev Genet 5, 725–738 (2004).

53. Davisson, M. T. et al. Segmental trisomy as a mouse model for Down syndrome. Prog Clin Biol Res 384, 117–133 (1993).

54. Reinholdt, L. G. et al. Molecular characterization of the translocation breakpoints in the Down syndrome mouse model Ts65Dn. Mamm Genome 22, 685–691 (2011).

55. Martinez, J. L., Zammit, M. D., West, N. R., Christian, B. T. & Bhattacharyya, A. Basal Forebrain Cholinergic Neurons: Linking Down Syndrome and Alzheimer’s Disease. Front Aging Neurosci 13, 703876 (2021).

56. Lindberg, M. F. et al. Chemical, Biochemical, Cellular, and Physiological Characterization of Leucettinib-21, a Down Syndrome and Alzheimer’s Disease Drug Candidate. J. Med. Chem. 66, 15648–15670 (2023).

57. Sonamoto, R. et al. Identification of a DYRK1A Inhibitor that Induces Degradation of the Target Kinase using Co-chaperone CDC37 fused with Luciferase nanoKAZ. Sci Rep 5, 12728 (2015).

58. Papenfuss, M. et al. Differential maturation and chaperone dependence of the paralogous protein kinases DYRK1A and DYRK1B. Sci Rep 12, 2393 (2022).

59. Belfiore, R. et al. Temporal and regional progression of Alzheimer’s disease-like pathology in 3xTg-AD mice. Aging Cell 18, e12873 (2019).

60. Tallino, S., Winslow, W., Bartholomew, S. K. & Velazquez, R. Temporal and brain region-specific elevations of soluble Amyloid-β40-42 in the Ts65Dn mouse model of Down syndrome and Alzheimer’s disease. Aging Cell 21, e13590 (2022).

61. Jones, B. J. & Roberts, D. J. The quantiative measurement of motor inco-ordination in naive mice using an acelerating rotarod. J Pharm Pharmacol 20, 302–304 (1968).

62. Dave, N. et al. Dietary choline intake is necessary to prevent systems-wide organ pathology and reduce Alzheimer’s disease hallmarks. Aging Cell 22, e13775 (2023).

63. Velazquez, R. et al. Acute tau knockdown in the hippocampus of adult mice causes learning and memory deficits. Aging Cell 17, e12775 (2018).

64. Alldred, M. J., Lee, S. H., Stutzmann, G. E. & Ginsberg, S. D. Oxidative Phosphorylation Is Dysregulated Within the Basocortical Circuit in a 6-month old Mouse Model of Down Syndrome and Alzheimer’s Disease. Front Aging Neurosci 13, 707950 (2021).

65. Houser, B. Bio-Rad’s Bio-Plex® suspension array system, xMAP technology overview. Arch Physiol Biochem 118, 192–196 (2012).

66. Teng, E. L., Chui, H. C., Schneider, L. S. & Metzger, L. E. Alzheimer’s dementia: performance on the Mini-Mental State Examination. J Consult Clin Psychol 55, 96–100 (1987).

67. Braak, H., Alafuzoff, I., Arzberger, T., Kretzschmar, H. & Del Tredici, K. Staging of Alzheimer disease-associated neurofibrillary pathology using paraffin sections and immunocytochemistry. Acta Neuropathol 112, 389–404 (2006).

68. Hyman, B. T. et al. National Institute on Aging–Alzheimer’s Association guidelines for the neuropathologic assessment of Alzheimer’s disease. Alzheimers Dement 8, 1–13 (2012).

69. Kinney, J. W. et al. Inflammation as a central mechanism in Alzheimer’s disease. Alzheimers Dement (N Y) 4, 575–590 (2018).

70. Plantone, D. et al. The Role of TNF-α in Alzheimer’s Disease: A Narrative Review. Cells 13, 54 (2023).

71. Karaman, M. W. et al. A quantitative analysis of kinase inhibitor selectivity. Nat Biotechnol 26, 127–132 (2008).

72. Becker, W. & Sippl, W. Activation, regulation, and inhibition of DYRK1A. The FEBS Journal 278, 246–256 (2011).

73. Himpel, S. et al. Identification of the autophosphorylation sites and characterization of their effects in the protein kinase DYRK1A. Biochem J 359, 497–505 (2001).

74. Dixon, A. S. et al. NanoLuc Complementation Reporter Optimized for Accurate Measurement of Protein Interactions in Cells. ACS Chem Biol 11, 400–408 (2016).

75. Branca, C. et al. Dyrk1 inhibition improves Alzheimer’s disease-like pathology. Aging Cell 16, 1146–1154 (2017).

76. Mattsson-Carlgren, N. et al. Longitudinal plasma p-tau217 is increased in early stages of Alzheimer’s disease. Brain 143, 3234–3241 (2020).

77. Suárez-Calvet, M. et al. Novel tau biomarkers phosphorylated at T181, T217 or T231 rise in the initial stages of the preclinical Alzheimer’s continuum when only subtle changes in Aβ pathology are detected. EMBO Molecular Medicine 12, e12921 (2020).

78. Regalado-Reyes, M. et al. Phospho-Tau Changes in the Human CA1 During Alzheimer’s Disease Progression. J Alzheimers Dis 69, 277–288 (2019).

79. Frost, D. et al. β-Carboline Compounds, Including Harmine, Inhibit DYRK1A and Tau Phosphorylation at Multiple Alzheimer’s Disease-Related Sites. PLOS ONE 6, e19264 (2011).

80. Simon, P., Dupuis, R. & Costentin, J. Thigmotaxis as an index of anxiety in mice. Influence of dopaminergic transmissions. Behav Brain Res 61, 59–64 (1994).

81. Fortea, J. et al. Down syndrome-associated Alzheimer’s disease: a genetic form of dementia. Lancet Neurol 20, 930–942 (2021).

82. Murphy, A. J., Wilton, S. D., Aung-Htut, M. T. & McIntosh, C. S. Down syndrome and DYRK1A overexpression: relationships and future therapeutic directions. Front Mol Neurosci 17, 1391564 (2024).

83. Kelley, C. M. et al. Sex differences in the cholinergic basal forebrain in the Ts65Dn mouse model of Down syndrome and Alzheimer’s disease. Brain Pathol 24, 33–44 (2014).

84. Tallino, S. et al. Assessing the Benefit of Dietary Choline Supplementation Throughout Adulthood in the Ts65Dn Mouse Model of Down Syndrome. Nutrients 16, 4167 (2024).

85. Hanson, S. M. et al. What makes a kinase promiscuous for inhibitors? Cell Chem Biol 26, 390–399.e5 (2019).

86. Liu, Q. et al. E3 Ligase SCFβTrCP-induced DYRK1A Protein Degradation Is Essential for Cell Cycle Progression in HEK293 Cells. J Biol Chem 291, 26399–26409 (2016).

87. Shukla, R., Kumar, A., Kelvin, D. J. & Singh, T. R. Disruption of DYRK1A-induced hyperphosphorylation of amyloid-beta and tau protein in Alzheimer’s disease: An integrative molecular modeling approach. Front. Mol. Biosci. 9, (2023).

88. Bilousova, T. et al. Synaptic Amyloid-β Oligomers Precede p-Tau and Differentiate High Pathology Control Cases. The American Journal of Pathology 186, 185–198 (2016).

89. Dubois, B. et al. Advancing research diagnostic criteria for Alzheimer’s disease: the IWG-2 criteria. Lancet Neurol 13, 614–629 (2014).

90. Hampel, H. et al. The Amyloid-β Pathway in Alzheimer’s Disease. Mol Psychiatry 26, 5481–5503 (2021).

91. Buchholz, S. & Zempel, H. The six brain-specific TAU isoforms and their role in Alzheimer’s disease and related neurodegenerative dementia syndromes. Alzheimer’s & Dementia 20, 3606–3628 (2024).

92. Yin, X. et al. Dyrk1A overexpression leads to increase of 3R-tau expression and cognitive deficits in Ts65Dn Down syndrome mice. Sci Rep 7, 619 (2017).

93. Jin, N. et al. Truncation and Activation of Dual Specificity Tyrosine Phosphorylation-regulated Kinase 1A by Calpain I: A MOLECULAR MECHANISM LINKED TO TAU PATHOLOGY IN ALZHEIMER DISEASE*. Journal of Biological Chemistry 290, 15219–15237 (2015).

94. Shi, J. et al. Increased dosage of Dyrk1A alters alternative splicing factor (ASF)-regulated alternative splicing of tau in Down syndrome. J Biol Chem 283, 28660–28669 (2008).

95. Woods, Y. L. et al. The kinase DYRK phosphorylates protein-synthesis initiation factor eIF2Bepsilon at Ser539 and the microtubule-associated protein tau at Thr212: potential role for DYRK as a glycogen synthase kinase 3-priming kinase. Biochem J 355, 609–615 (2001).

96. Glick, D., Barth, S. & Macleod, K. F. Autophagy: cellular and molecular mechanisms. J Pathol 221, 3–12 (2010).

97. Congdon, E. E. et al. Methylthioninium chloride (methylene blue) induces autophagy and attenuates tauopathy in vitro and in vivo. Autophagy 8, 609–622 (2012).

98. Hamano, T. et al. Autophagic-lysosomal perturbation enhances tau aggregation in transfectants with induced wild-type tau expression. Eur J Neurosci 27, 1119–1130 (2008).

99. Ozcelik, S. et al. Rapamycin Attenuates the Progression of Tau Pathology in P301S Tau Transgenic Mice. PLoS One 8, e62459 (2013).

100. Wang, P. et al. DYRK1A Interacts with the Tuberous Sclerosis Complex and Promotes mTORC1 Activity. eLife 12, (2023).

101. Liu A et al. MicroRNA-215-5p inhibits the proliferation of keratinocytes and alleviates psoriasis-like inflammation by negatively regulating DYRK1A and its downstream signalling pathways. Experimental dermatology 30, (2021).

102. Khor, B. et al. The kinase DYRK1A reciprocally regulates the differentiation of Th17 and regulatory T cells. eLife 4, e05920 (2015).

103. Tarkowski, E., Andreasen, N., Tarkowski, A. & Blennow, K. Intrathecal inflammation precedes development of Alzheimer’s disease. J Neurol Neurosurg Psychiatry 74, 1200–1205 (2003).

104. Appleton, J. et al. Brain inflammation co-localizes highly with tau in mild cognitive impairment due to early-onset Alzheimer’s disease. Brain awae234 (2024) doi:10.1093/brain/awae234.

105. Chen, Y. & Yu, Y. Tau and neuroinflammation in Alzheimer’s disease: interplay mechanisms and clinical translation. Journal of Neuroinflammation 20, 165 (2023).

106. Kitazawa, M. et al. Blocking IL-1 signaling rescues cognition, attenuates tau pathology, and restores neuronal β-catenin pathway function in an Alzheimer’s disease model. J Immunol 187, 6539–6549 (2011).

107. Leyns, C. E. G. & Holtzman, D. M. Glial contributions to neurodegeneration in tauopathies. Mol Neurodegener 12, 50 (2017).

108. Parbo, P. et al. Does inflammation precede tau aggregation in early Alzheimer’s disease? A PET study. Neurobiol Dis 117, 211–216 (2018).

109. Bhaskar, K. et al. Regulation of Tau Pathology by the Microglial Fractalkine Receptor. Neuron 68, 19–31 (2010).

110. Ben Menachem-Zidon, O., Menahem, Y. B., Hur, T. B. & Yirmiya, R. Intra-Hippocampal Transplantation of Neural Precursor Cells with Transgenic Over-Expression of IL-1 Receptor Antagonist Rescues Memory and Neurogenesis Impairments in an Alzheimer’s Disease Model. Neuropsychopharmacology 39, 401–414 (2014).

111. Zhang, J., Ke, K.-F., Liu, Z., Qiu, Y.-H. & Peng, Y.-P. Th17 Cell-Mediated Neuroinflammation Is Involved in Neurodegeneration of Aβ1-42-Induced Alzheimer’s Disease Model Rats. PLOS ONE 8, e75786 (2013).

112. Rueda, N. et al. Anti-IL17 treatment ameliorates Down syndrome phenotypes in mice. Brain Behav Immun 73, 235–251 (2018).

